# Mechanistic insight into light-dependent recognition of Timeless by Drosophila cryptochrome

**DOI:** 10.1101/2021.09.10.459772

**Authors:** Changfan Lin, Connor M. Schneps, Siddarth Chandrasekaran, Abir Ganguly, Brian R. Crane

## Abstract

Cryptochrome (CRY) entrains the fly circadian clock by binding to Timeless (TIM) in light and triggering its degradation. Undocking of a helical C-terminal tail (CTT) in response to photoreduction of the CRY flavin cofactor gates TIM binding. A generally-applicable Select Western-blot-Free Tagged-protein Interaction (SWFTI) assay enables quantification of CRY binding to TIM in dark and light. The assay is utilized to study CRY variants with residue substitutions in the flavin pocket and correlate their TIM affinities with CTT undocking, as measured by pulse-dipolar ESR spectroscopy and evaluated by molecular dynamics simulations. CRY variants with the CTT removed or undocked bind TIM constitutively, whereas those incapable of photoreduction bind TIM weakly. In response to flavin redox state, two conserved histidine residues contribute to a robust on/off switch by mediating CTT interactions with the flavin pocket and TIM. Our approach provides an expeditious means to quantify protein-protein interactions and photoreceptor targeting.

## INTRODUCTION

Cryptochromes (CRYs) belong to a light-sensitive flavoprotein superfamily that includes the DNA repair enzyme photolyase (Conrad et al., 2014; Crane and Young, 2014; Sancar, 2003). CRYs play important roles in circadian clock regulation (Figure 1A) and are classified into two types according to their functions: Type I CRYs serve as photoreceptors to reset circadian rhythms in animals and plants, whereas type II CRYs act as light-independent transcriptional oscillators in mammals (Conrad *et al.*, 2014; Fogle et al., 2015; Ozturk, 2017). In the model clock of fruit flies (*Drosophila melanogaster*) type I CRY has been shown to have multiple functions, both as a photoreceptor in key pacemaker neurons, and as transcription regulators in circadian oscillators of peripheral cells (Fogle *et al.*, 2015; Foley and Emery, 2020).

**Figure 1:**
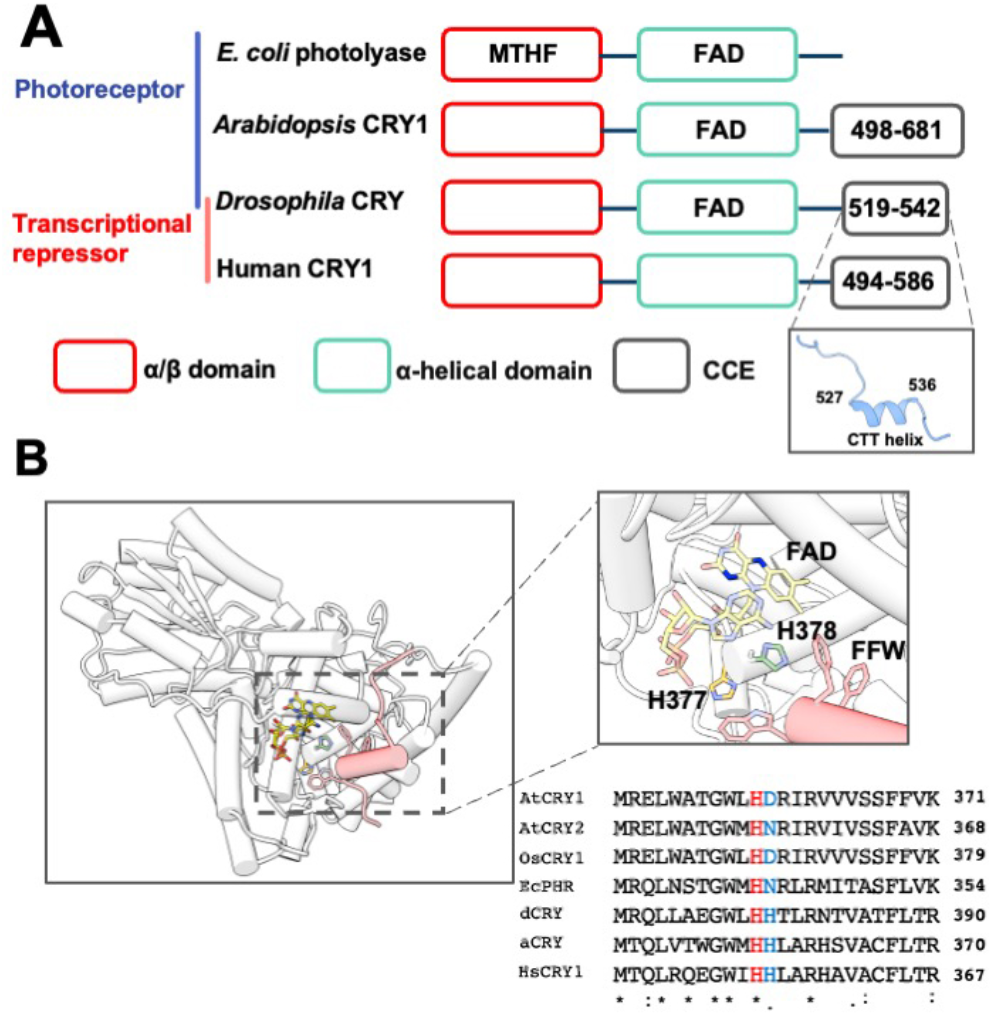
The flavin-pocket and CTT of Drosophila cryptochrome (dCRY) (A) Region maps of cryptochromes and *E. coli* photolyases. The proteins share a photolyase homology region (PHR) comprising a conserved α/β domain (red) that binds the antenna molecule MTHF in photolyases and α-helical domain (green) that contains the FAD-binding pocket. Cryptochromes have a Cryptochrome C-terminal Extension (CCE, gray) of variable lengths. In dCRY, the CCE forms a CTT helix structure. *E. coli* photolyase and Type I cryptochrome Arabidopsis CRY1 are photoreceptive, whereas Type II cryptochromes such as human CRY1 function as transcriptional repressors in the circadian clock; dCRY may have both roles in flies. (B) Residues His377 (orange, with nitrogen colored blue) and His378 (green, with nitrogen colored blue) are located between cofactor FAD (colored by heteroatoms, nitrogen, light blue; oxygen, red; phosphorus, orange; N1 and N5 of the isoalloxazine ring which are involved in reduction reactions, dark blue) and FFW motif (Phe534, Phe535, Trp536, red) of the CCE (red, helices shown as cylinders) in CRY (PDB: 4GU5). The amino acid sequence of CRY in this region is aligned with other cryptochrome/photolyase proteins, such as *Arabidopsis thaliana* CRY1/2 (AtCRY1/2), *E. coli* photolyase (EcPHR), *Homo sapiens* CRY1 (HsCRY1)*, Chlamydomonas reinhardtii animal-like cryptochrome* (aCRY), *and Oryuza sativa japonica* CRY1 (OsCRY1).

In *Drosophila*’s central nervous system, the 24 hr clock cycle is maintained by a transcriptional translational feedback loop (TTFL) involving the proteins Timeless (TIM) and Period (PER). TIM and PER form heterodimers in the cytosol, enter the nucleus, and bind to the transcription factors Clock and Cycle, opposing their enhancer activities and thus inhibiting the production of TIM, PER and other clock-controlled proteins. This molecular machinery is entrained to the environment by light-activated CRY that recruits the E3 ligase Jetlag (JET) to TIM for proteasome-based degradation of the PER:TIM complex (Koh et al., 2006; Peschel et al., 2009). CRY itself also undergoes light-triggered degradation by JET and/or another E3 ligase known as Ramshackle (BRWD3) (Ozturk et al., 2013).

It remains of great interest how light promotes CRY-TIM heterodimerization. All CRYs have a variable CRY C-terminal Extension (CCE) that appends the conserved photolyase homology region (PHR) and appears to regulate their function (Figure 1A) (Chaves et al., 2011; Emery et al., 1998; Levy et al., 2013; Ozturk et al., 2007; Zoltowski et al., 2011). In *Drosophila* CRYs, the CCE takes the form of a C-terminal tail helix (CTT) that binds into the flavin pocket analogously to how damaged DNA substrates are recognized by photolyase (Levy *et al.*, 2013; Zoltowski *et al.*, 2011). Removal of the CTT results in constitutive binding of CRY to TIM (Dissel et al., 2004) and accelerates TIM degradation (Busza et al., 2004). A number of studies have established that the CTT releases during light activation (Berntsson et al., 2019; Chandrasekaran et al., 2021; Ganguly et al., 2016; Lin et al., 2018; Ozturk et al., 2014). Most likely the release is triggered by photoreduction of the flavin to the anionic semiquinone state, of which the negative charge on the isoalloxazine ring is necessary to promote the conformational change (Chandrasekaran *et al.*, 2021; Ganguly *et al.*, 2016; Lin *et al.*, 2018). Molecular dynamics (MD) simulations, residue substitution studies and the pH dependence of photoreduction kinetics suggested that coupled protonation of a conserved His378, which resides between the flavin and the Phe534-Phe535-Trp536 (FFW) motif of the CTT, disrupted interactions between the CTT and the flavin pocket (Figure 1B). Substitutions of His378 to Asn, Arg, or Lys, alter CTT conformation and TIM or CRY degradation behavior in complex ways (Chandrasekaran *et al.*, 2021; Ganguly *et al.*, 2016). However, time-resolved small-angle x-ray scattering (SAXS) experiments have shown that the H378A variant behaves similarly to WT with respect to light-induced global conformational changes monitored by SAXS (Berntsson *et al.*, 2019).

To better understand the mechanism of CTT gating, one would ideally like to quantify changes in the binding affinity between CRY and TIM in dark and light. However, such measurements are hampered by the fact that TIM is a large, post-translationally modified protein, with regions of low complexity and intrinsic disorder that make it challenging to produce and purify. Thus, gold standard methods to quantify protein-protein interactions such as isothermal titration calorimetry, fluorescent polarization assays, surface plasmon resonance, and biointerferometry are generally not possible (Rao et al., 2014; Syafrizayanti et al., 2014). Historically, yeast-two hybrid (Y2H) assays have been used to identify the interaction between TIM and CRY variants or JET and their light-dependence (Hemsley et al., 2007; Peschel *et al.*, 2009). However, Y2H is generally not quantitative and post-translational modifications and cofactor availability may be different in yeast than in *Drosophila* cells. Besides owing to a propensity for false positives, further validation is often required. Immunoprecipitation coupled with western blotting has also been used to study the CRY:TIM complex with either CRY or TIM as the bait (Koh *et al.*, 2006; Peschel *et al.*, 2009). However, unreliable quantification of protein amounts in western blots is common, resulting, for example, from unpredictable sample loss during blot transfer. In addition, western blotting involves tedious wash steps and indirect chemiluminescent detection that can detract from the ability to accurately correlate protein abundance with the enzyme activity of conjugated antibodies (Butler et al., 2019; Ghosh et al., 2014; Pillai-Kastoori et al., 2020). In the case of monitoring CRY activity, we and others (Ganguly *et al.*, 2016; Ozturk *et al.*, 2014) have relied on monitoring decreases in TIM levels with light; however, such measurements reflect many other cellular processes and rely on the assumption that CRY binding to TIM is the limiting factor.

Thus, to better characterize the impact of key residue substitutions on CTT release we developed a binding assay that relies on the fluorescent detection of tagged proteins that can be expressed directly in insect cells. We then coupled this functional assay with an electron-spin-resonance (ESR)-spectroscopy spin-labeling approach that directly monitors CRY CTT release (Chandrasekaran *et al.*, 2021). These two methods provide insight into the role of two conserved His residues in the flavin binding pocket and reveal a critical CRY residue involved in not only CTT undocking but also TIM recognition.

## RESULTS

### A western-blot free pull-down assay to quantify CRY-TIM interactions

We introduce a Select Western-blot-Free Tagged-protein Interaction (SWFTI) assay based on a SNAP/CLIP-tag based fluorescent detection method to quantify CRY-TIM complexation in Drosophila S2 cells, eliminating the need to purify TIM. SNAP/CLIP tags are protein domains of only 20 kDa (182 residues), even smaller than the common fluorescent protein tag GFP (28 kDa), and thus they minimally interfere with protein structure and function (Gautier et al., 2008; Keppler et al., 2003). The SNAP tag covalently attaches to any fluorophore conjugated with benzylguanine, whereas the complementary CLIP tag enables an orthogonal reaction with a benzylcytosine derivative (Figure 2A). Compared to fluorescent proteins, fluorophores are more photostable and they remain associated with proteins of interest, even under denaturing conditions, such as boiling and SDS-PAGE.

**Figure 2:**
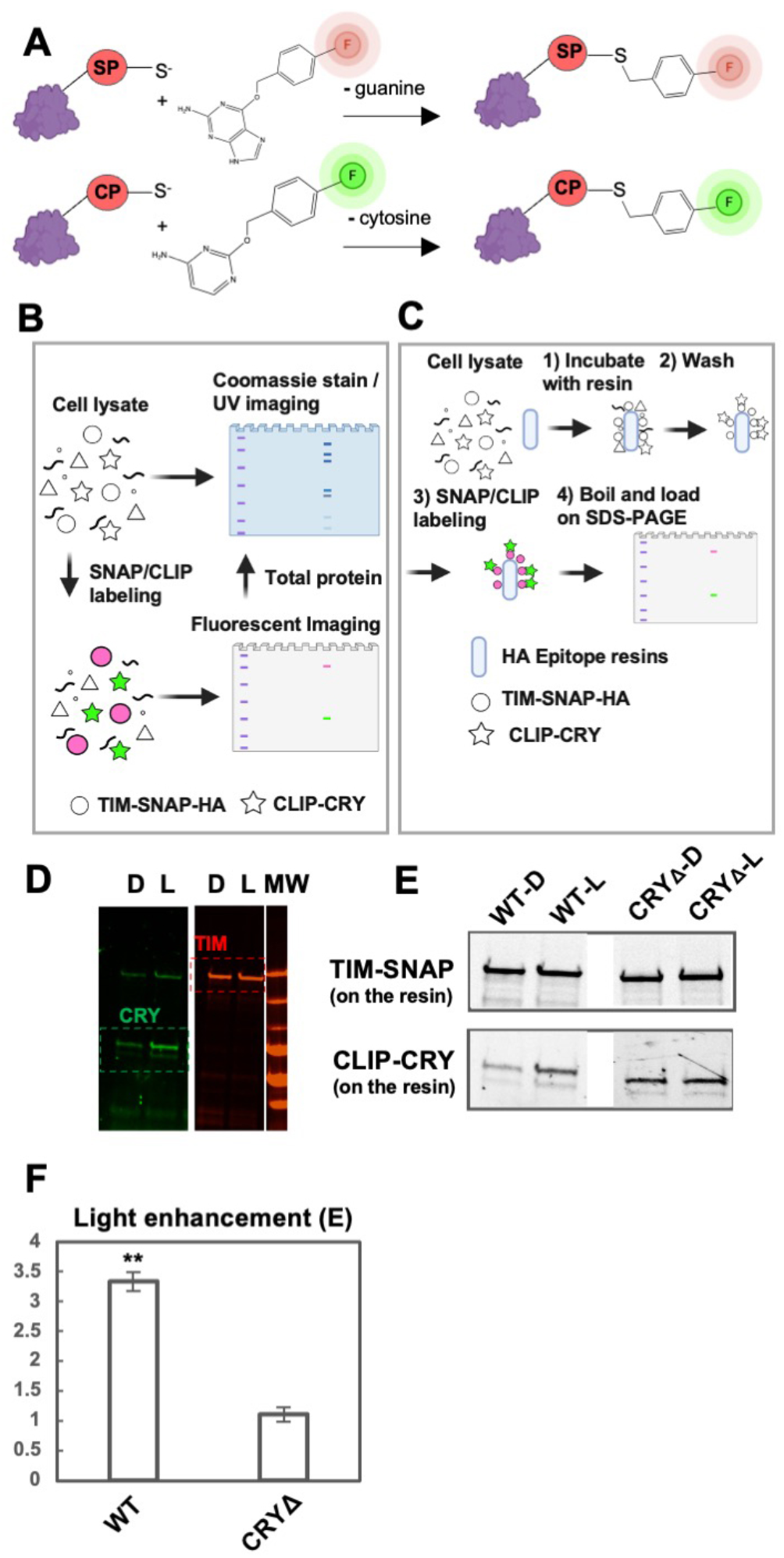
The SWFTI assay (A) Orthogonal reactions of the SNAP/CLIP tags that conjugate benzylguanine/benzylcytosine fluorescent reporter moieties to the protein tag (red, green). (B) CRY variants and the TIM proteins were engineered with SNAP/CLIP fusion tags, shown as TIM-SNAP-HA (circles) and CLIP-CRY (stars). The fusion proteins are specifically labeled in cell lysate with different fluorophores simultaneously: benzylguanine in green and benzylcytosine in pink. Detection is achieved by multiplex fluorescent imaging of a resolving gel (SDS-PAGE or native gel; TIM: red band, CRY: green band, ladder: purple), whereas the total protein is monitored by Coomassie stain or UV-induced imaging (BioRad, stain-free gel). Symbols made with biorender.com. (C) For pull-downs: 1) TIM-CRY complex in cell lysate is enriched by HA epitope resins using TIM-SNAP-HA as the bait. 2) HA resin was washed with TBS-T buffer to remove non-specific binding. 3) TIM-SNAP-HA and CLIP-CRY react directly with SNAP/CLIP fluorophores on resin. 4) Resin was boiled and denatured in SDS protein sample buffer to elute fluorescent labeled TIM and CRY. Protein amount was evaluated by SDS-PAGE (TIM: red band, CRY: green band). (D) Representative SWFTI results for CRY-wild type (86 kDa, green) and TIM (179 kDa, red) on HA resin. CRY displayed enhanced interaction with TIM in the light-exposed sample (L) compared with the dark sample (D). Both dark and light-exposed samples are aliquoted from the same culture dish to ensure the same protein amount in the lysate. MW = molecular weight markers. (E-F) CRY-wild type (WT, residue 1-539) exhibited light enhanced affinity with TIM, whereas CRYΔ (residue 1-520) showed no dark/light difference. Light enhancement is defined as E= (CRY on resin / TIM on resin)_light_ / (CRY on resin / TIM on resin)_dark_. Error bars reflect the SEM for n = 3. A two-tailed single sample t-Test was performed with a hypothetical mean of 1, thus a significant difference from 1 indicates light-dependent interaction between CRY variants and TIM. ** p value ≤ 0.01.

Owing to robust reaction conditions, we were able to achieve multiplex labeling and imaging of both TIM and CRY in either lysate or on purification resins (Figures 2B and 2C). We constructed CLIP-CRY and TIM-SNAP-HA fusion genes in an S2 cell expression vector and transfected them into S2 cells for transient expression. To capture CRY-TIM complexes but prevent subsequent TIM degradation due to endogenous JET in S2 cells (Peschel *et al.*, 2009), the proteasome inhibitor MG132 was added to transfected cells prior to light exposure. SNAP and CLIP dyes with compatible fluorescence spectra were added directly to the cell lysate, the lysate was loaded on SDS-PAGE gels, and the resulting gels were imaged in a multiplexed format (Figure 2B). Total protein amounts on the same gel can also be monitored by Coomassie staining or UV light-induced stain-free imaging (Gilda and Gomes, 2013). To probe CRY-TIM interactions, cells were harvested and lysed after illumination, and CRY-TIM complexes were enriched by HA-antibody resin using TIM-SNAP-HA as bait (Figure 2C). The relative amounts of CRY and TIM bound to the resin were analyzed with a multi-channel fluorescent imager after SDS-PAGE. Fluorescence intensities relative to an internal standard (CLIP-CRY-SNAP construct) gave the relative amount of TIM and CRY bound to the resin. The fraction of TIM bound to CRY (*F*) was calculated using equation (1), based on a 1:1 binding stoichiometry between TIM and CRYΔ (which comprises residues 1-520 and does not contain the CTT). The binding stoichiometry was verified to be 1.08 by quantifying the relative components of the complex with the blot-free method under non-denaturing conditions (Figure S1).

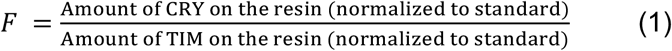

The light-dependence of CRY-TIM interaction was then evaluated by light enhancement (*E*), which is defined as equation (2); an *E* value of 1 indicates no enhancement of binding in the light.

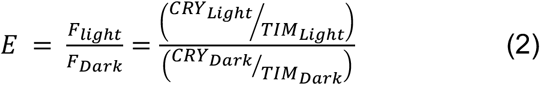

With this method, we investigated TIM binding by CRY-wild type (WT) and CRYΔ, which has been previously shown to bind TIM more strongly in the dark (Busza *et al.*, 2004; Peschel *et al.*, 2009). The HA resin pulled down similar amounts of TIM bait in each sample (Figures 2D-E). WT exhibited nearly 3.5 × more CRY binding in light, whereas CRYΔ showed tight light-independent TIM binding (Figures 2F, 3).

### Relative dark versus light binding affinities

Changes to light enhancement (*E*; equation 2) observed with different CRY variants does not distinguish whether the causative residue substitutions affect dark-state binding, light-state binding or both. Lower *E* values could result from an increase in dark-state affinity or a reduction in light-state affinity for TIM. For example, CRYΔ shows no light/dark discrimination (*E* ~ 1) because with the CTT removed there is no gating; i.e. CRYΔ binds TIM strongly in both dark and light. Thus, we sought a method to evaluate the binding affinity between CRY and TIM. Although not an equilibrium condition, pull-down reactions can be used to estimate an effective dissociation constant (K_D_) under the assumption that the resin largely captures the equilibrium binding distribution prior to pull-down and the dissociation time constants are not much faster than seconds. K_D_ is then calculated based on equations below (Equation 3–5). F is determined from equation (1).

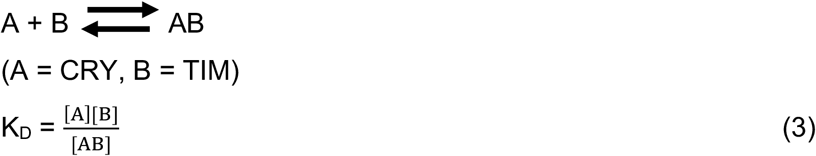

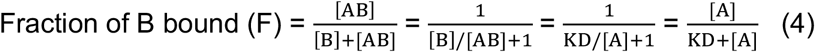

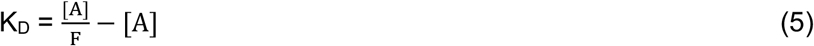

In our experiments [A] >> [AB] (Figure S1), therefore [*A*] ≈ [*A*] + [*AB*] = *A*_*total*_, where *A*_*total*_ is the sum of bound and unbound *A* in the cell lysate. We used purified SNAP proteins of known concentrations to quantify the concentration of the CLIP-CRY-SNAP standard, which is then used to calculate absolute values of [*A*] under identical conditions of illumination (Figure S2). Once the binding affinity of the dark sample (K_D,dark_) has been determined, that of the light sample (K_D,light_) can be derived from this value and *E* (Equation 6).

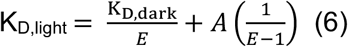

Under these conditions, WT CRY binds TIM with a K_D,dark_ of ~ 32 μM. K_D,light_ decreases to 9 μM (Figure 3B,C). Although the redox potential of CRY is quite low (−320 mV) (Lin *et al.*, 2018), when overexpressed, some CRY may be reduced, which would then cause an underestimation of K_D,dark_ however, we would expect this effect to be similar for all of the variants. CRYΔ, shows a high, light-insensitive affinity for TIM (K_D,dark_ = 1.7 μM; K_D,light_ = 1.5 μM), which indicates that the docked CTT greatly reduces TIM binding and that the undocked CTT does not contribute substantially to TIM binding. Unlike CRYΔ, which binds to TIM strongly in dark and light, The W394F variant, which does not undergo FAD photoreduction (Lin *et al.*, 2018), displays weak binding to TIM regardless of light condition (Figure 3C). This behavior is then reversed with removal of the CTT in variant W394FΔ (residues 1-520; Figure 3C). Thus, photoreduction of FAD causes CTT undocking, which allows TIM binding.

**Figure 3:**
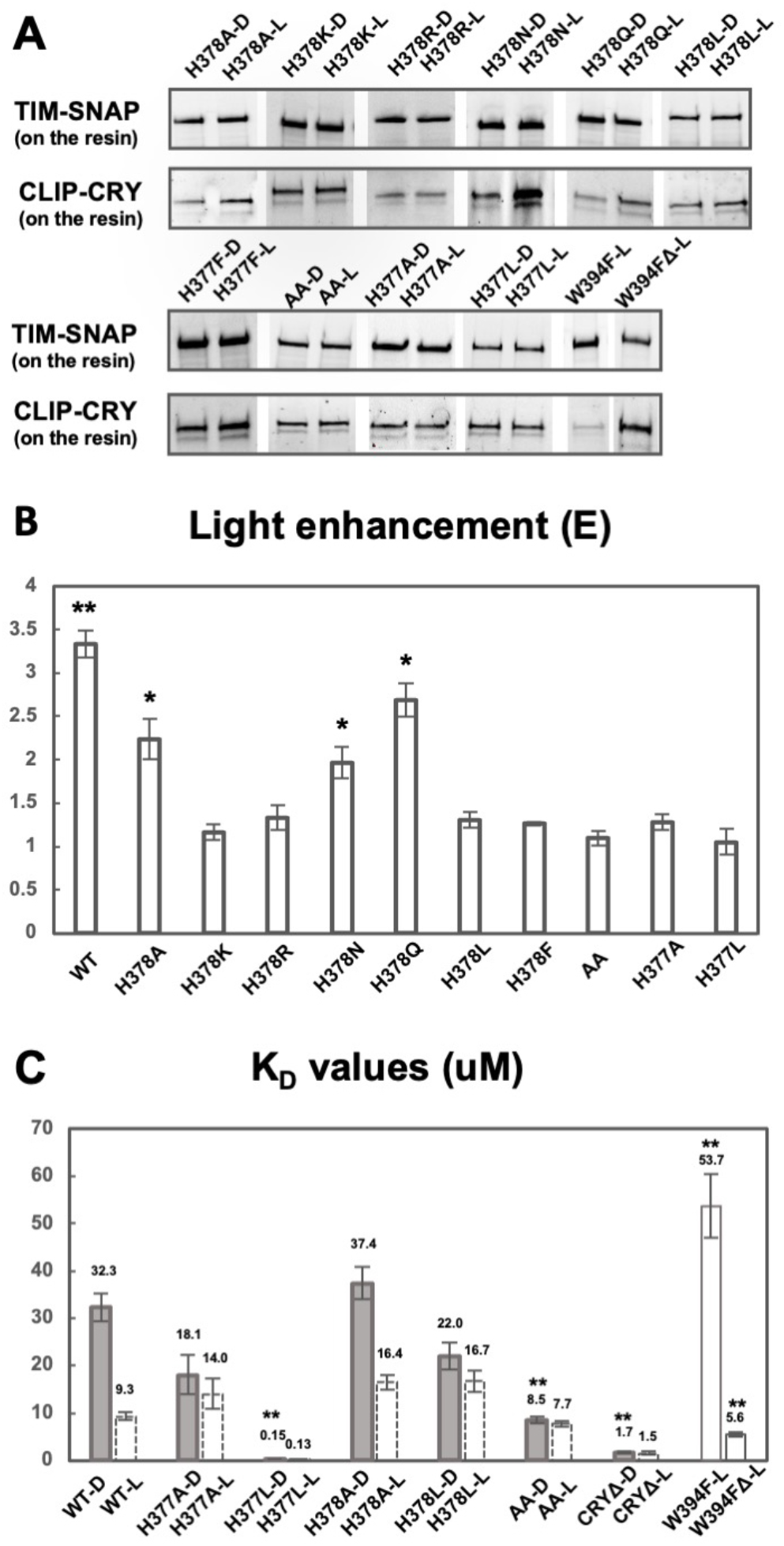
His377 and His378 are key residues for gating the CRY-TIM interaction (A) SWFTI assessment of binding affinities of TIM for CRY variant in dark (D) and in light (L). W394FΔ represents W394F mutant of CRYΔ (residue 1-520). AA represents the H377A/H378A CRY variant. Data is representative of 3 individual replicates, except that H378F has 2 replicates. (B) Quantification of 3A. Light-enhancement defined as in Figure 2E. A two-tailed single sample t-Test was performed with a hypothetical mean of 1. *p value ≤ 0.05. **p value ≤ 0.01. (C) K_D_ values reflect the relative binding strength between CRY variants and TIM with tighter interactions giving a smaller K_D_ values. Dashed columns indicate values calculated based on the K_D_ of corresponding dark samples and the light enhancements shown Figures 2F and 3B. All solid columns were compared to WT-D by one-way ANOVA. * Tukey HSD p value ≤0.05, ** Tukey HSD p value ≤ 0.01.

### Changes to conserved His residues alter TIM binding in light and dark

CRY contains two His residues in the vicinity of the CTT, His377 and His378. His378 is conserved in type I CRYs and His377 is nearly invariant across all CRY proteins (Figure 1B); both residues are also found in photolyases. His377 interacts with Trp536, producing an edge on contact between the two respective aromatic rings, but projects away from the flavin pocket. In contrast, His378 resides between the isoalloxazine ring and the FFW motif of the CTT. MD simulations suggest that the N1P tautomer (i.e. Nδ protonated) of the neutral His378 imidazole stabilizes the CTT against the PHR with the oxidized or reduced flavin, but that a doubly protonated imidazole or an N3P tautomer (i.e. Nε protonated) promotes tail undocking, the latter only in the case of the reduced flavin (Ganguly *et al.*, 2016). We have also shown that the His378Asn (N) partially releases the CTT in the dark and degrades TIM prior to light exposure (Ganguly *et al.*, 2016). Interestingly the His378Arg (R), His378Lys (K) variants allow relatively normal TIM degradation, but completely block light-dependent degradation of CRY itself, while displaying both some undocking in the dark state and incomplete release in the light state (Chandrasekaran *et al.*, 2021; Ganguly *et al.*, 2016). Residues involved in altering the CTT conformation may also interface with TIM directly, and thus we sought methods to directly quantify both CTT undocking and TIM binding in the variants.

If the protonation state of H378 is important for its hydrogen bonding network with the CTT, then replacing it with K/R or N/Gln (Q), which have hydrogen bonding properties not affected by protonation at physiological pH, should alter light-dependent TIM interactions. As expected, H378K/R exhibited no significant light-dependence to the TIM interaction whereas H378N and H378Q still retained light-enhanced binding of TIM, although to a reduced extent compared to His378 (Figure 3A, B). H378Q gave near WT behavior, which perhaps correlates with it being a steric analog of the His N3P tautomer. H378A does discriminate light from dark, but to a reduced amplitude compared to WT (Figure 3B). We also examined His378Leu (L) and His378Phe (F), which have a size similar to His, but also no side-chain hydrogen bonding capacity, and found low dark-light contrast in both variants. Thus, H378K/R/F/L variants do not show light/dark discrimination for TIM. In contrast, the H378N/Q variants bind TIM better in light, although the switch has lower fidelity than does WT. Hence, a protonation change at residue 378 is not a requirement for CTT undocking, but residues incapable of undergoing a change in protonation state generally show little light-enhanced binding of TIM.

Protonation changes at His377 may compensate for loss of protonation changes when His378 is substituted. Even though His377 does not lie directly between the flavin and the CTT, the His377 side chain contacts Trp536 and resides near the flavin pocket. Consistent with a role for His377 in the CTT gating mechanism, both the H377A and the H377A/H378A double substitution gave no light enhancement of TIM binding (Fig. 3A, B).

### H377A/H378A exhibits tighter dark-state binding to TIM than WT and H378A

Examination of the relative dissociation constants indicates that the His377/His378 Ala or Leu variants show a range of behaviors, but most increase dark-state TIM affinity, with the double variant H377A/H378A (AA) binding to TIM in both the dark and light as well as WT binds to TIM in the light (Figure 3C). AA shows little light sensitivity, binds to TIM strongly in the dark, but in neither dark nor light binds TIM as tightly as CRYΔ. H378A gives near WT behavior in the dark with a reduced affinity in the light, whereas H377A has increased dark-state affinity (K_D_ = 18.1 μM) and a light-state affinity similar to H378A. Substitution of the His residues to the hydrophobic Leu residue, which is of similar size, has a greater impact than the Ala variants. H378L closely resembles H377A in TIM binding behavior, giving an increased dark-state affinity and a slightly reduced light-state affinity. However, H377L produces dramatic effects both in dark and light, binding to TIM in both conditions with an affinity that is ~10x greater than even CRYΔ (Figure 3C).

### Increased TIM binding in the dark correlates with CTT undocking

To probe the effects of His377 and His378 on CTT conformational stability and dynamics, we utilized CTT specific spin-labeling via sortylation and a L405E/C416N (“EN”) CRY variant that forms a flavin neutral semiquinone (NSQ), but does not release the CTT in the otherwise WT protein (Figure 4A) (Chandrasekaran *et al.*, 2021). The neutral flavin radical of the EN variant serves as a dark-state proxy that can be used as a reference to measure the distance distribution of a spin-label on the C-terminus with pulse-dipolar ESR spectroscopy (PDS). For the light-state, the flavin anionic semiquinone (ASQ) serves as the reference point. Alanine substitutions were made in both the wild-type-like (H377A, H378A, H377A/H378A) and dark-state proxy (H377A/L405E/C416N, H378A/L405E/C416N, H377A/H378A/L405E/C416N) proteins to elucidate the proportions of docked and undocked states. To ensure that the substitutions to His377 and His378 did not alter the redox states of the flavin, UV-Vis and cw-ESR spectra of all variants were taken in the dark and light. As expected, the single and double substitutions (H377A, H378A, H377A/H378A) all formed the anionic semiquinone (ASQ), whereas the triple and quadruple substitutions (H377A/L405E/C416N, H378A/L405E/C416N, H377A/H378A/L405E/C416N) formed the neutral semiquinone (NSQ) (Figures S3–S4). 4-pulse double electron-electron resonance (4P-DEER) spectroscopy was then performed on these variants, and the relative proportions of docked and undocked states were determined using several analysis procedures (see methods) as reported previously (Chandrasekaran *et al.*, 2021). Representing the resulting distance distributions as a sum of Gaussian functions by the program DD (Stein et al., 2015) describe the time domain traces relatively well under the assumption of a simple two-state model with the components restrained to the known WT distances (Figures 4B, S5). Unrestrained fits gave similar results (Figures S6,S7 and Table S2). As a secondary approach, singular value decomposition (SVD) resolved additional components and improved the fits to the time domain data (Figure S8). The SVD distance distributions agree qualitatively with those produced by the Gaussian fits (Figure S9). As an additional check, all time domain traces of the His377 and His378 variants were fit directly with linear combinations of the WT and L405E/C416N time domain traces; quantitatively, the percentages of each time domain component in the reconstructions agree well with those calculated by Gaussian two-state models (Figure S10).

**Figure 4:**
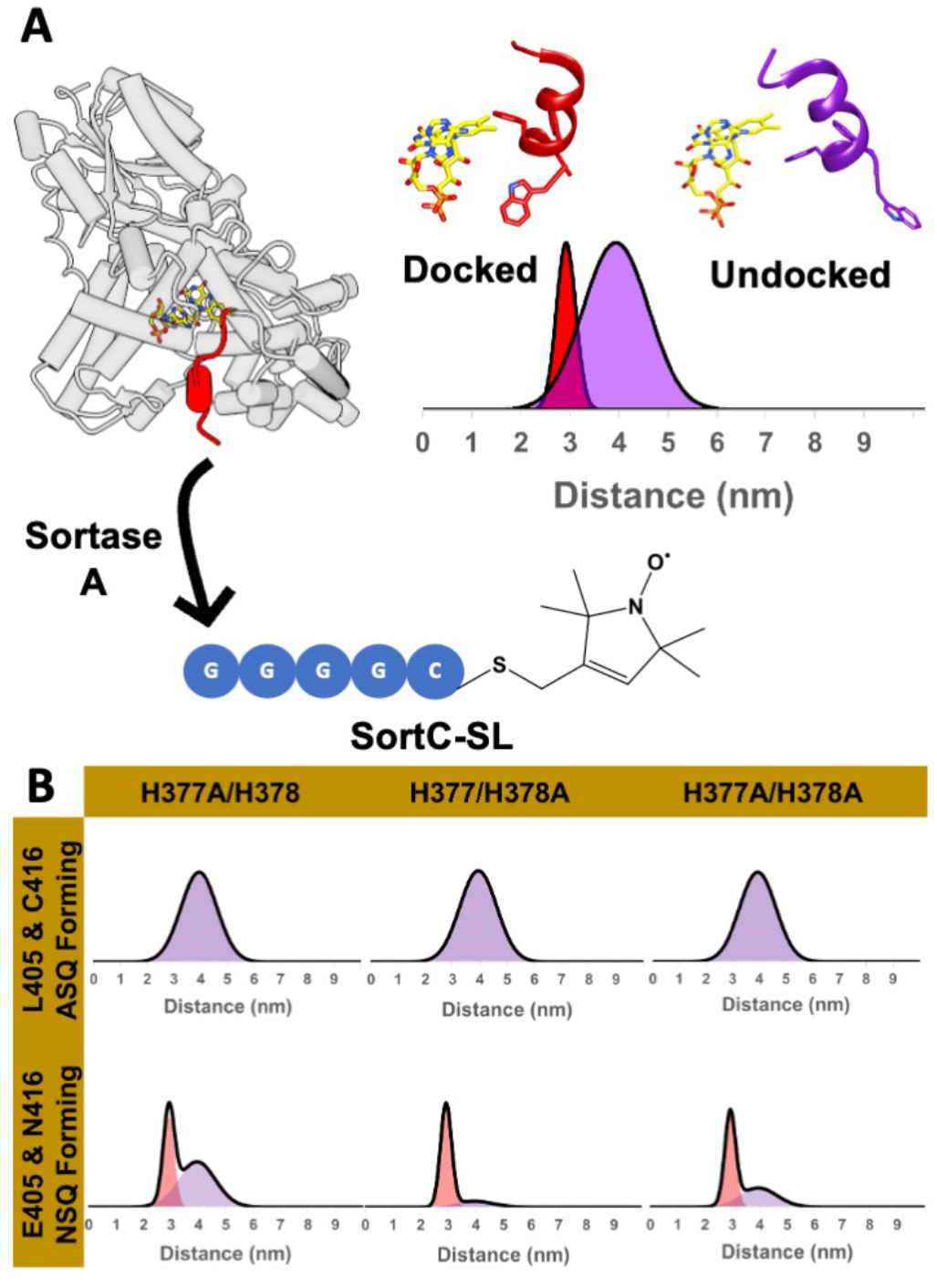
His377 and His378 stabilize the CTT against the flavin pocket in the dark. (A) PDS-based spin-spin distance distributions between a flavin radical and a C-terminal nitroxide moiety appended to the CTT (red) by site-enzymatic ligation of a spin-labeled peptide. (B) Distance distributions obtained after restraining to a one or two-state model of Gaussian function. ASQ forming variants were restrained to one-component (the WT parent, green), while the NSQ forming variants (purple) were restrained to two components (mixture of docked/undocked states).

DEER spectroscopy of the H378A variant reveals that it is wild-type like in the PDS assay, in that the CTT is fully undocked in the light state, while a large percentage of it is docked in the dark-state proxy (Figure 4B). Like H378A, the H377A variant exhibits WT-like CTT undocking in the light, but unlike H378A, H377A shows a proportion of undocking in the dark-state proxy. Taken together, the data imply that His377 plays a more important role in stabilizing the CTT against the flavin pocket in the dark than does His378, and neither residue on their own is required for undocking. DEER data on the double variant H377A/H378A (AA) both in the WT and EN background indicates a similar distance distribution in the light compared to WT; however, in the EN background the AA retains a higher proportion of the docked state than does H377A alone, suggesting that His378 destabilizes the CTT in the absence of interactions otherwise provided by His377 in the NSQ state. Given that H377L binds TIM more strongly than CRYΔ in the dark we examined the CCT conformation of H377L in the EN background (H377L/L405E/C416N). H377L EN, much like H377A, produces a substantial proportion of the undocked state (Figure S11). However, H377L binds TIM much more tightly in the dark than H377A, and even more than CRYΔ Thus, in addition to affecting CTT dynamics Leu377 likely directly forms interactions within the TIM interface.

### Molecular Dynamics Simulations

Following previous work (Chandrasekaran *et al.*, 2021; Ganguly *et al.*, 2016), we examined the conformational stability of the CTT in the His377/His378 variants with MD simulations. Consistent with those studies, the CTT is stable against the flavin pocket when the flavin is in its oxidized form and the His residues contain neutral imidazole moieties (Figure 5A). The H377A substitution gives a relatively stable CTT with the oxidized flavin, and similar stability with the NSQ (Figure 5B), which is consistent with the binding data (Figure 3C). However, in the presence of the ASQ the CTT displaces, even when His378 is neutral and contains the N1P tautomer (Figure 5B). In contrast, the AA variant shows a relatively unstable CTT in the presence of oxidized as well as ASQ flavin. For both AA and H377A, the CTT appears more mobile in the NSQ state than with the oxidized flavin (Figure 5B,C), which would suggest that the NSQ state only approximates the oxidized flavin state when residue substitutions perturb interactions of the CTT.

**Figure 5:**
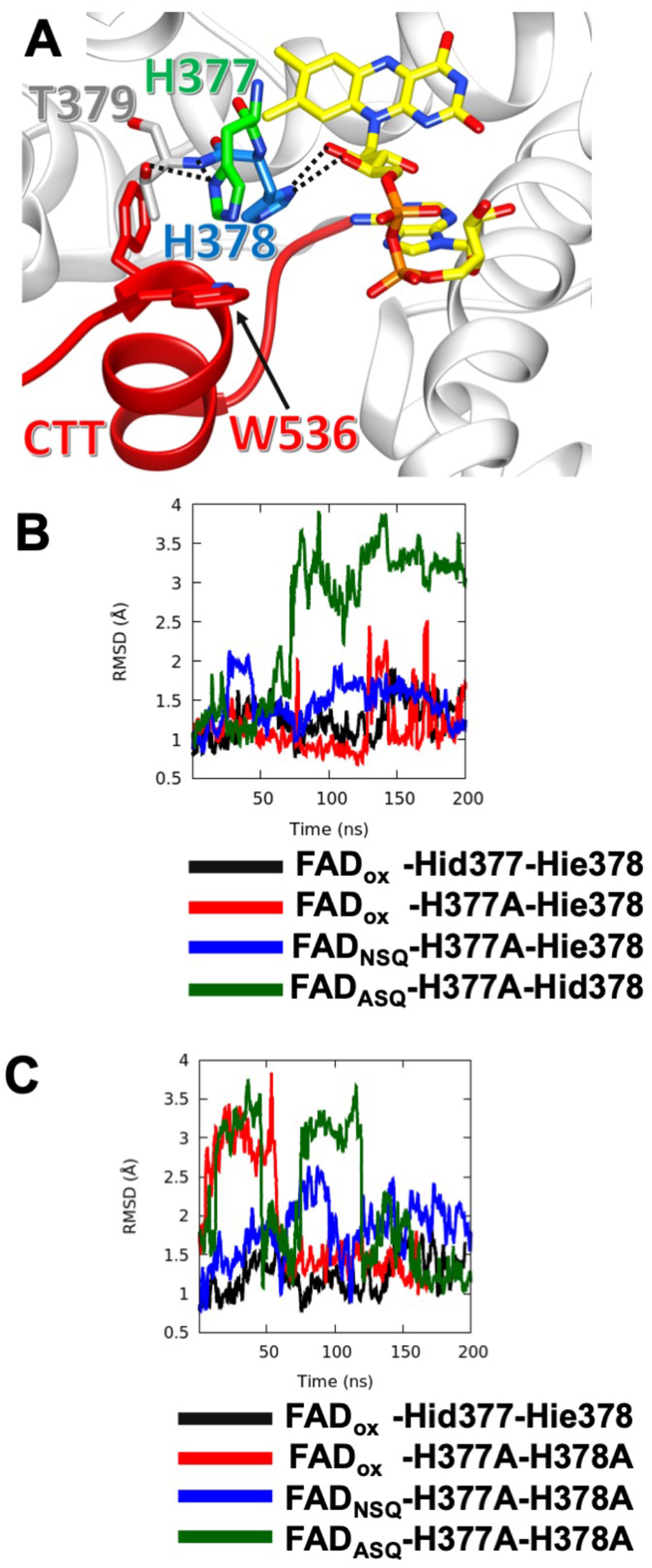
MD Simulations reveal varying stabilities of the CTT in CRY variants. (A) Local bonding environment of His377 and His378. His377 (green) makes hydrogen bonding contacts with the amide nitrogen of the backbone and the hydroxyl group of Thr379; His377 also engages in edge-on van der Waal’s interactions with the sidechain of Trp536 of the FFW (red).

## DISCUSSION

Structure function studies of CRY have been hampered by the lack of quantitative assays for the light-dependent interaction between CRY and TIM that underlies entrainment of the circadian clock. Such methods have been challenging to develop largely because TIM is difficult to procure outside of cells. The SWFTI assay provides fluorescent quantification of interacting proteins and absolute determination of protein amounts with relatively fast and efficient readouts. Furthermore, covalent attachment of the detection dyes to each fusion tag renders the assay insensitive to protein denaturation as required by SDS-PAGE separation. The SWFTI method should be widely useful for the quantification of protein-protein interactions in a cellular context. To showcase this assay, we demonstrate how FAD photoreduction and the CTT conformation impact the affinity of CRY for TIM and investigate the role that two conserved His residues play in modulating interactions between the CRY CTT and the PHR.

The CRY CTT is known to act as an autoinhibitory element for TIM interactions (Busza *et al.*, 2004; Dissel *et al.*, 2004). Photoreduction of the CRY flavin to the ASQ causes CTT undocking (Lin *et al.*, 2018; Vaidya et al., 2013). In our experiments where both proteins are overexpressed within insect cells, WT CRY binds TIM to some extent in the dark, but the measured affinity increases substantially in the light. Residue substitutions that lie in the interface between the CTT and the flavin pocket alter the dark state affinity of CRY by destabilizing the bound conformation of the CTT. Consistent with this interpretation, CRYΔ shows high affinity for TIM in both light and dark and no enhanced binding in light. Similarly, the W394F variant that cannot be photoreduced and does not release the CTT (Lin *et al.*, 2018) shows only weak affinity for TIM in dark and light. Changes in light state affinity of CRY for TIM then likely owe to a partially released CTT, and/or changes (direct or indirect) in the CRY:TIM interaction surface resulting from the substitutions.

Photoactive proteins, both natural and designed, typically change affinity for their targets by at least an order of magnitude when in light-activated states (Akiyama et al., 2016; Kepsutlu et al., 2014; Kondoh and Terazima, 2017; Zimmerman et al., 2016). Whereas a dark state affinity in the ~30 μM range for CRY is likely sufficient to arrest TIM degradation, an increase in affinity for TIM by only ~3.5-fold for the WT protein seems modest. However, CRYΔ binds to TIM more tightly than light-state CRY (with a K_D_ in the 1 μM range), indicating that for the WT protein, the lower light-state affinity may be due to incomplete light-activation. A contributing factor to incomplete conversion may be that the pH environment of the cell converts some of the flavin ASQ to the NSQ (Einholz et al., 2021), thereby reducing the amount of CRY bound to TIM in light. Furthermore, some apo CRY during overexpression (which we estimate could be as high as 25~) will not contribute to TIM binding but will register in the free sample. Dark state affinity may also be overestimated for several reasons. Although the reduction potential of the CRY flavin is low (Lin *et al.*, 2018), some chemically reduced CRY may increase dark-state TIM binding. Moreover, the equilibrium between CTT docking/undocking, though favoring the docked state in the dark, may produce enough undocked CTT to bind some TIM under conditions where CRY is excess. It is difficult to distinguish among these possibilities except perhaps to note that CRY does not show any reduced semiquinone on purification from insect cells. The comparison of K_D_ values between CRYΔ and W394F perhaps give the best indication of the true switch in affinity upon activation (35-fold difference) and indicate that in these conditions WT CRY in the dark is somewhat overactivated, whereas WT CRY in the light is somewhat underactivated. Finally, it should be emphasized that although these assays provide accurate relative measures of affinity, the absolute values are estimates. CRY may dissociate during the pull-down and wash steps (owing to flavin reoxidation, for example) and only a few regions of the binding isotherm are sampled in these experiments owing to the achievable cellular expression levels. If the actual dissociation constants are far off the concentrations of free CRY and TIM in the cells, the quantification will be inaccurate. In that sense, it is reassuring that we are able to observe a > 2 order of magnitude difference in effective binding constants among the variants.

In consideration of the CTT release mechanism, His378 has drawn attention because of its position between the flavin and the FFW motif, its conservation in Type I cryptochromes and the known change in protonation state analogous His residues undergo in the photolyase mechanism (Berntsson *et al.*, 2019; Ganguly *et al.*, 2016; Hitomi et al., 2001; Schleicher et al., 2007). MD simulations indicate that His378 protonation destabilizes the CTT against the flavin pocket but that the neutral N3P tautomer also undocks the CTT when the flavin is reduced to the ASQ. Furthermore, substitution of His378 removes a modest pH dependence to photoreduction rates (Ganguly *et al.*, 2016). However, time-resolved SAXS experiments suggest that the H378A variant behaved very similarly to WT with respect to light-induced conformational changes (Berntsson *et al.*, 2019). Indeed, our interaction data showed that the H378A protein binds TIM in both light and dark similar to WT, but with a lower light enhancement. Based on our PDS assays, H378A resembles WT in CTT release behavior, with slightly more population of the undocked CTT in the EN dark-state proxy. However, this increased proportion of the undocked state in H378A is not reflected by increased affinity for TIM in the dark and H378A instead shows reduced affinity in the light. These changes in TIM recognition suggest that although H378A approximates WT-like behavior the fidelity of the CTT conformational switch has been disrupted to some extent. Nearby His377 could also undergo protonation changes when the flavin reduces to the ASQ, either as a primary site, or as a secondary site when His378 has been replaced by a non-ionizable residue. His377 hydrogen bonds to the amide nitrogen of Thr379 and is thereby likely neutral in the resting oxidized state of the protein (Figure 5A). Changing His377 to Ala increases TIM binding somewhat in the dark, and little there is little enhancement of binding in the light. This data would suggest that the CTT is likely somewhat destabilized by the H377A substitution and conversion to the undocked state has also been curtailed. The PDS data on the other hand indicates a more drastically destabilized CTT in both dark and light. The double H377A/H378A variant (AA) binds TIM in dark and light much tighter than WT, approaching the affinity of CRYΔ. By PDS, AA shows less undocking in the dark than H377A, which does not align with the trend in the affinity data. This result perhaps reflects the limitation of the NSQ as a dark-state proxy. The MD simulations bear out a conformationally destabilized CTT in the AA variant and suggest greater stability of the CTT for H377A, which is consistent with the binding affinity data.

Whether flavin reduction promotes protonation of His377 or His378 is still unclear because most substitutions alter both CTT conformation in the dark as well as light enhancement of TIM binding. Taking TIM binding as a measure of CTT release, substitution of either His residue to various other residues does not block CTT release completely, but light-state affinity for TIM is often reduced relative to WT (e.g. H377A, H378A,L), and light enhanced binding is lost in cases where the CTT is still appreciably bound in the dark-state proxy (e.g. H378K,R). The AA variant also loses light enhancement entirely, but the high affinity of TIM binding in the dark indicates that the substitutions perturb the dark-state CTT conformation and thereby largely remove gating.

An additional complication is that residues surrounding the CTT likely play a role in directly binding TIM. For example, the H377L substitution undocks the CTT in the dark state proxy to the same extent as the AA (Figure S9), but TIM binding is extremely strong to this variant in both dark and light. Because the affinity is an order of magnitude higher than even CRYΔ, the effect must involve more than just CTT dynamics: TIM most likely binds to this region and Leu377 stabilizes this interaction. Consistent with this idea, a peptide derived from TIM that resembles the CTT sequence preferentially pulls down CRY in the light (Vaidya *et al.*, 2013). The ability of H378R,K to perturb the CTT to varying extents, but produce no light enhancement of TIM binding also suggests that substitutions at this position interfere with TIM binding. As noted previously, H378R,K stabilized CRY against ubiquitin mediated degradation (Ganguly *et al.*, 2016), thereby suggesting that the E3-ligase machinery may also be directed toward this region of the flavin pocket.

Overall, our data indicates 1) that TIM binding requires CTT undocking and the distribution of docked and undocked conformations correlate with TIM affinity; 2) flavin reduction is required for CTT undocking and the resulting enhancement of TIM binding is of reasonable magnitude to regulate the system; 3) the undocking mechanism is sensitive to the 377 and 378 residues, with the highest fidelity switching requiring His at both positions and 4) TIM very likely interfaces with the pocket exposed by CTT undocking. The general role of CCE in gating and modulating the interactions of different CRY proteins with both targets and small molecules is well established and has been recently discussed in detail (Chandrasekaran *et al.*, 2021; Crane, 2020; Miller et al., 2020b; a; Miller et al., 2020c; Parico and Partch, 2020). Although in many cases the molecular details remain to be worked out, the root of the sensing mechanisms involve light- or ligand-induced shifts in equilibria of docked and undocked CCEs, which thereby control binding to CRY interactors. Recent MD simulations of CRY photoactivation suggest that disruption of an Asp-Arg salt bridge next to the flavin ring rapidly follows flavin reduction and couples to CTT release (Wang et al., 2021). The tools utilized here, provide a means to assay and track these processes in a cellular context, correlate them with biophysical properties and thereby test and refine our understanding of the underlying molecular mechanisms.

**Table 1.** Fitting statistics for restrained Gaussian Model. Residual χv^2^ values are determined by DD and indicate the robustness of the fit to the data. Values below 2 indicate a good fit. H377A, H378A, and AA are all WT-like and were fit best using one-component assuming complete undocking (*). These data have a relatively high error (**), but resemble the SVD reconstructions of Figure S9 qualitatively.

| Variant | Radical | % Undocked | $\chi_v^2$ (error in fit) |
| --- | --- | --- | --- |
| H377A | ASQ | 100* | 3.1** |
| H378A | ASQ | 100* | 7.1** |
| H377A/H378A | ASQ | 100* | 1.4 |
| H377A/L405E/C416N | NSQ | $63 \pm 1$ | 1.4 |
| H378A/L405E/C416N | NSQ | $17 \pm 0.3$ | 24.** |
| H377A/H378A/L405E/C416N | NSQ | $42 \pm 2$ | 1.0 |

While His378 hydrogen bonds to the ribosyl oxygens of the FAD. (B) The running average of CTT displacement with respect to the crystal conformation (closed state) is monitored over the course of various MD trajectories. Deviations in CTT conformation are observed to varying extents with the H377A variant, depending on flavin redox state. FADox – CRY with an oxidized isoalloxazine ring; NSQ – the flavin neutral semi-quinone; ASQ – the flavin anionic semiquinone; Hid – His with N1(δ) protonated; Hie – His with N3(ε) protonated; Hip – His with both N1(δ) and N3(ε) protonated. CTT conformational stability shown relative to Hid377/Hie378 for comparison (black line). The running average is calculated over a window of 400 ps. (C) MD simulations of CTT displacements in the H377A/H378A variant as a function of flavin redox state. Nomenclature as in (B).

## ACKNOWLEDGMENTS

This work was financially supported by NIH grant R35GM122535 (B.R.C.). ESR measurements were carried out at ACERT which is supported by NIH/NIGMS awards P41 GM103521 and 1S1 0OD021543. Sortase A plasmid was a gift from William DeGrado at UCSF.

## AUTHOR CONTRIBUTIONS

C.L., C.M.S., and B.R.C. designed research; and C.L., C.M.S., A.G. and B.R.C. performed research; C.L., C.M.S., A.G., S.C. and B.R.C. analyzed data; C.L., C.M.S., and B.R.C. wrote the manuscript with contributions from all authors.

## Materials and Methods

### DNA sequences and constructs

1) CLIP-CRY variants (in pAC5.1 vector)

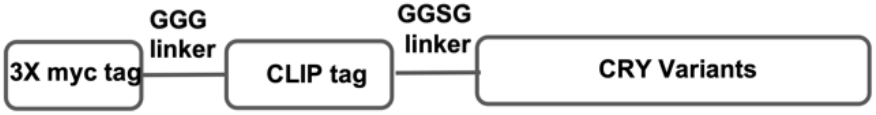

2) TIM-SNAP-HA (in pAC5.1 vector)

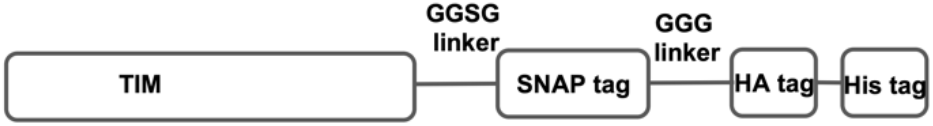

3) CLIP-CRY-SNAP (in pAC5.1 vector)

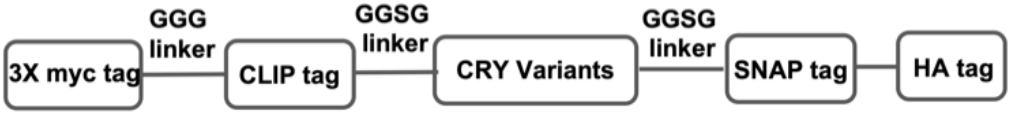

4) CRY variants with sort-tag (in pET28 vector)

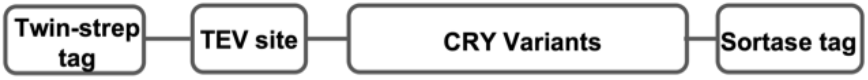

The following *Drosophila* specific kozak sequence was used to initiate protein translation in pAC5.1 vectors:

### AAAAATGG

#### CRY-WT (1-539)

ATGGCCACGCGAGGGGCGAATGTGATTTGGTTTCGCCATGGATTGCGCCTCCATGATAATCCCGCTCTATTGGCCGC CCTCGCCGATAAGGATCAGGGTATAGCCCTAATTCCCGTTTTCATATTCGATGGAGAGAGTGCAGGTACCAAGAATG TGGGTTACAATCGGATGCGTTTCCTCCTGGACTCGTTGCAGGACATCGATGATCAGCTACAGGCGGCAACTGATGGA CGTGGACGCCTCCTGGTCTTCGAGGGCGAACCGGCTTATATCTTCCGCCGGCTACATGAGCAAGTGCGTCTGCACAG GATTTGCATAGAGCAGGACTGCGAGCCAATTTGGAATGAGCGCGATGAAAGCATCCGTTCTCTATGTCGGGAGCTGA ATATCGACTTTGTCGAGAAGGTATCACACACGCTTTGGGATCCGCAATTGGTGATTGAGACCAATGGTGGCATTCCA CCGCTGACCTACCAAATGTTCCTGCACACGGTGCAAATTATTGGGCTTCCACCGCGTCCCACCGCCGATGCTCGACT AGAAGACGCCACCTTTGTCGAGCTGGACCCCGAGTTCTGCCGAAGTCTTAAGTTGTTCGAGCAGCTGCCCACGCCGG AGCACTTCAATGTGTATGGAGACAACATGGGCTTCCTGGCCAAGATTAACTGGCGCGGCGGAGAAACACAGGCCTTA CTTCTGTTGGATGAGCGTCTTAAAGTGGAGCAGCATGCGTTTGAGCGTGGATTTTATCTGCCCAACCAGGCACTGCC CAATATCCACGACTCGCCAAAATCGATGAGCGCCCATCTGCGCTTTGGTTGCCTTTCGGTACGTCGCTTCTACTGGA GCGTCCACGATCTCTTCAAGAATGTCCAGTTGCGCGCCTGTGTGCGGGGCGTTCAGATGACTGGCGGCGCGCACATC ACGGGACAGTTGATCTGGCGAGAGTACTTCTACACCATGTCGGTGAACAATCCAAACTACGATCGCATGGAGGGCAA TGACATCTGCCTGAGCATCCCGTGGGCTAAGCCGAACGAAAATCTCCTGCAGAGCTGGCGTTTAGGCCAAACGGGAT TCCCGCTCATCGACGGCGCCATGCGACAACTCCTGGCCGAGGGATGGCTCCACCATACGCTGCGCAACACCGTGGCC ACCTTTCTCACGCGCGGCGGTTTGTGGCAGAGCTGGGAGCATGGACTGCAGCACTTTCTGAAGTATCTGCTGGATGC GGATTGGTCGGTCTGCGCTGGCAACTGGATGTGGGTATCCAGCTCGGCGTTTGAAAGGCTGCTGGACTCCTCCCTGG TCACCTGCCCGGTGGCATTGGCCAAGCGACTTGATCCGGATGGCACCTACATCAAGCAGTACGTCCCGGAGTTGATG AATGTGCCCAAGGAATTTGTTCACGAGCCCTGGCGAATGTCTGCCGAGCAGCAGGAGCAGTACGAGTGCCTGATCGG AGTCCATTATCCGGAGCGGATCATTGATTTGTCCATGGCCGTAAAGCGAAACATGCTGGCCATGAAGTCTCTCCGAA ATTCGCTGATCACCCCGCCGCCGCATTGCCGACCATCCAACGAGGAGGAAGTGCGTCAGTTCTTCTGGCTGGCCGAC

#### CRYΔ (1-520)

ATGGCCACGCGAGGGGCGAATGTGATTTGGTTTCGCCATGGATTGCGCCTCCATGATAATCCCGCTCTATTGGCCGCCCTCGCCGATAAGGATCAGGGTATAGCCCTAATTCCCGTTTTCATATTCGATGGAGAGAGTGCAGGTACCAAGAATGTGGGTTACAATCGGATGCGTTTCCTCCTGGACTCGTTGCAGGACATCGATGATCAGCTACAGGCGGCAACTGATGGACGTGGACGCCTCCTGGTCTTCGAGGGCGAACCGGCTTATATCTTCCGCCGGCTACATGAGCAAGTGCGTCTGCACAGGATTTGCATAGAGCAGGACTGCGAGCCAATTTGGAATGAGCGCGATGAAAGCATCCGTTCTCTATGTCGGGAGCTGAATATCGACTTTGTCGAGAAGGTATCACACACGCTTTGGGATCCGCAATTGGTGATTGAGACCAATGGTGGCATTCCACCGCTGACCTACCAAATGTTCCTGCACACGGTGCAAATTATTGGGCTTCCACCGCGTCCCACCGCCGATGCTCGACTAGAAGACGCCACCTTTGTCGAGCTGGACCCCGAGTTCTGCCGAAGTCTTAAGTTGTTCGAGCAGCTGCCCACGCCGGAGCACTTCAATGTGTATGGAGACAACATGGGCTTCCTGGCCAAGATTAACTGGCGCGGCGGAGAAACACAGGCCTTACTTCTGTTGGATGAGCGTCTTAAAGTGGAGCAGCATGCGTTTGAGCGTGGATTTTATCTGCCCAACCAGGCACTGCCCAATATCCACGACTCGCCAAAATCGATGAGCGCCCATCTGCGCTTTGGTTGCCTTTCGGTACGTCGCTTCTACTGGAGCGTCCACGATCTCTTCAAGAATGTCCAGTTGCGCGCCTGTGTGCGGGGCGTTCAGATGACTGGCGGCGCGCACATCACGGGACAGTTGATCTGGCGAGAGTACTTCTACACCATGTCGGTGAACAATCCAAACTACGATCGCATGGAGGGCAATGACATCTGCCTGAGCATCCCGTGGGCTAAGCCGAACGAAAATCTCCTGCAGAGCTGGCGTTTAGGCCAAACGGGATTCCCGCTCATCGACGGCGCCATGCGACAACTCCTGGCCGAGGGATGGCTCCACCATACGCTGCGCAACACCGTGGCCACCTTTCTCACGCGCGGCGGTTTGTGGCAGAGCTGGGAGCATGGACTGCAGCACTTTCTGAAGTATCTGCTGGATGCGGATTGGTCGGTCTGCGCTGGCAACTGGATGTGGGTATCCAGCTCGGCGTTTGAAAGGCTGCTGGACTCCTCCCTGGTCACCTGCCCGGTGGCATTGGCCAAGCGACTTGATCCGGATGGCACCTACATCAAGCAGTACGTCCCGGAGTTGATGAATGTGCCCAAGGAATTTGTTCACGAGCCCTGGCGAATGTCTGCCGAGCAGCAGGAGCAGTACGAGTGCCTGATCGGAGTCCATTATCCGGAGCGGATCATTGATTTGTCCATGGCCGTAAAGCGAAACATGCTGGCCATGAAGTCTCTCCGAAATTCGCTGATCACCCCGCCG

### 3x Myc tags

**GAGCAGAAGCTGATCTCAGAGGAGGACCTG**GGAGGAGGA**GAACAAAAATTAATAAGTGAAGAAGACCTG**GGCGGCGG C**GAGCAGAAGCTGATCTCAGAGGAGGACCTG**

#### CLIP tag

ATGGACAAAGACTGCGAAATGAAGCGCACCACCCTGGATAGCCCTCTGGGCAAGCTGGAACTGTCTGGGTGCGAACAGGGCCTGCACCGTATCATCTTCCTGGGCAAAGGAACATCTGCCGCCGACGCCGTGGAAGTGCCTGCCCCAGCCGCCGTGCTGGGCGGACCAGAGCCACTGATCCAGGCCACCGCCTGGCTCAACGCCTACTTTCACCAGCCTGAGGCCATCGAGGAGTTCCCTGTGCCAGCCCTGCACCACCCAGTGTTCCAGCAGGAGAGCTTTACCCGCCAGGTGCTGTGGAAACTGCTGAAAGTGGTGAAGTTCGGAGAGGTCATCAGCGAGAGCCACCTGGCCGCCCTGGTGGGCAATCCCGCCGCCACCGCCGCCGTGAACACCGCCCTGGACGGAAATCCCGTGCCCATTCTGATCCCCTGCCACCGGGTGGTGCAGGGCGACAGCGACGTGGGGCCCTACCTGGGCGGGCTCGCCGTGAAAGAGTGGCTGCTGGCCCACGAGGGCCACAGACTGGGCAAGCCTGGGCTGGGT

#### SNAP tag

ATGGACAAAGACTGCGAAATGAAGCGCACCACCCTGGATAGCCCTCTGGGCAAGCTGGAACTGTCTGGGTGCGAACAGGGCCTGCACCGTATCATCTTCCTGGGCAAAGGAACATCTGCCGCCGACGCCGTGGAAGTGCCTGCCCCAGCCGCCGTGCTGGGCGGACCAGAGCCACTGATGCAGGCCACCGCCTGGCTCAACGCCTACTTTCACCAGCCTGAGGCCATCGAGGAGTTCCCTGTGCCAGCCCTGCACCACCCAGTGTTCCAGCAGGAGAGCTTTACCCGCCAGGTGCTGTGGAAACTGCT GAAAGTGGTGAAGTTCGGAGAGGTCATCAGCTACAGCCACCTGGCCGCCCTGGCCGGCAATCCCGCCGCCACCGCCGCCGTGAAAACCGCCCTGAGCGGAAATCCCGTGCCCATTCTGATCCCCTGCCACCGGGTGGTGCAGGGCGACCTGGACGTGGGGGGCTACGAGGGCGGGCTCGCCGTGAAAGAGTGGCTGCTGGCCCACGAGGGCCACAGACTGGGCAAGCCTGGGCTGGGT

#### HA tag

TACCCATACGATGTTCCAGATTACGCT

#### His tag

CATCATCACCATCACCAT

#### Twin-strep tag

TGGAGCCATCCGCAGTTTGAAAAAGGTGGTGGTAGCGGTGGTGGTTCAGGTGGTAGTGCATGGAGCCACCCGCAGTTCGAAAAA

#### Sortase tag

CTGCCGGGTACGGGC

#### TEV site

GAAAACCTGTACTTCCAATCC

### Gibson assembly

All constructs in pAC5.1 and pET28 vectors were built using the modified Gibson assembly method (Gibson et al., 2009). The master mix was prepared as described in Table S1, aliquoted to 15 μL each tube and stored at −20 °C. The insert and pAC5.1 vector were amplified by Q5 DNA polymerase-based PCR using primers designed by NEBuilder (New England Biolabs), with the standard parameters changed to provide 30 bp overlaps at the junction of the fragments. 50 μL PCR products were incubated with 1 μL DpnI enzyme (New England Biolabs) to clean up templates at 37 °C for 2.5 hours and then purified by gel-extraction following the manufacturer instructions (Cat# D2500-02, OMEGA Bio-tek). Then 50 to 100 ng purified vector was added to 15 μL of the Gibson master mix prior to adding purified insert to the mix at a molar ratio of 3:1 insert per vector (a ratio of 7:1 was used when the insert was less than 500 bp). The total volume was taken to 20 μL with nanopure water and the mixture was incubated at 50 °C for 15 to 30 minutes. After the reaction, 10 μL reaction products were added to 100 μL DH5α competent cells for transformation. The cells were incubated on ice for 30 minutes before heat shock at 42 °C for 45 seconds. Then 1 mL sterile LB media was added to the transformation tube without any antibiotics and shaken at 37 °C at 250 rpm for 1 hour. The mix was pelleted at 13,000 × g for 2 minutes at room temperature to concentrate the cell density. 800 μL supernatant was removed from the tube and the cell pellets were resuspended in the remaining ~200 μL supernatant and then spread on LB-agar plates for incubation overnight at 37 °C. Resulting colonies were processed following miniprep kit instruction (Cas# D6942-02, OMEGA Bio-tek) and sequenced at Cornell Institute of Biotechnology.

**Table S1:** Components of Gibson assembly master mix. Reagents all purchased from New England Biolabs, except PEG-8000 (Cat# HR21-535, Hampton Research) and NAD^+^ (Cat# 16077, Cayman Chemical).

| | Stock concentration (U/μL) | Final Concentration (U/μL) | Volume (μL) |
| --- | --- | --- | --- |
| Q5 reaction buffer | 5 X | 1.3 X | 234 |
| Q5 DNA polymerase | 2 | 0.033 | 14.8 |
| Taq DNA ligase | 40 | 5.3 | 119.2 |
| T5 Exonuclease (dilute with 1X NEbuffer 4) | 0.5 | 0.005 | 9 |
| PEG-8000 | 50% | 6.7% | 120.6 |
| NAD<sup>+</sup> | 50 mM (33.2 mg/ml) | 1.3 mM | 23.4 |
| dNTPs | 2.5 mM | 0.27 mM | 97.2 |
| Water |  |  | 282 |
| Total |  |  | 900 |

### Q5 DNA polymerase-based mutagenesis

To produce CRY variants, modified Q5 DNA polymerase-based mutagenesis was used. Briefly, primers were designed using CRY-WT-CLIP as the template by NEBaseChanger (New England Biolabs). The desired DNA was amplified by PCR as above. PCR products were purified by a gel extraction kit and then 100 ng purified products were mixed with 1 μL each of T4 DNA ligase buffer (10x), T4 DNA ligase, T4 Polynucleotide Kinase and DpnI (New England Biolabs), to heal nicks in the PCR products. Nanopure water was added to make the total volume 10 μL. After overnight incubation at room temperature, all 10 μL wwere used for transformation following the same protocol as above.

### Maintenance of S2 insect cells

Drosophila S2 cells were purchased from ATCC (Cat# CRL-1963). Cells were grown at 27 degrees in Schneider’s Drosophila Medium (Cat# 21720024, ThermoFisher) suppled with 10% heat-inactivated Fetal Bovine Serum (FBS, Cat# F4135, sigma). Cells were seeded at a density of 100 × 10^4^ cells/ml, and subcultured every 2 to 3 days when the cell density reached ~ 1000 × 10^4^ cells/ml.

### Transient transfection of S2 cells

pAC5.1 vectors with CLIP-CRY and TIM-SNAP-HA were delivered into s2 cells using Effectene reagent (Cat# 301425, Qiagen) or TransIT-Insect Transfection Reagent (Cat# MIR6100, Mirus) following modified manufacturer instruction. Briefly, 1000×10^4^ cells in 10 mL growth media with FBS were placed in a 100 mm culture dish (Cat# 353003, Corning) on day 1. About 20 hours later, 4 μg DNA in total (CRY variants: TIM mass ratio 1:1, except CRYΔ: TIM = 7:1) was used with Effectene transfection reagent or 10 μg DNA when using TransIT-Insect reagent (same CRY: TIM mass ratio as using Effectene). 72 hours after transfection, 50 μM (S)-MG132 (Cat# 10012628, Cayman Chemical) was added to the 10 mL cells. After a brief mixing period, 10 mL cells were evenly split into two 60 mm culture dishes (Cat# 353002, Corning) to ensure the same cell population for dark and light conditions. After 2 hours of incubation, light sample was placed on a LED light pad (Autograph PRO1200) with light filter sheets to only pass light with wavelength of 440 nm to 500 nm. Then cells were centrifuged at 500 ×g for 5 minutes, room temperature, washed once with chilled PBS and stored at −80 °C until lysis.

### The SWFTI assay

To lyse S2 cells, 400 μL lysis buffer with gentle non-ionic detergent (50 mM Tris buffer, pH 8, 150 mM NaCl, 1% IGEPAL CA-630 (Cat# I8896, Sigma Aldrich), 10% glycerol and 1x protease inhibitor (Cat# A32965, ThermoFisher)) was added to cell pellets harvested from 5 mL cells. EDTA was omitted to avoid disturbing SNAP/CLIP dye labeling. Note all buffer pH should be adjusted at the working temperature. After cells were incubated with lysis buffer on ice for 30 mins (flicking the tube every 10 min to avoid vortexing), cell debris was removed by spinning at 15,000 × g for 15 minutes at 4 °C.

To label the lysate sample with SNAP/CLIP dyes, 1 part of lysate was diluted with 2 parts of labeling buffer (50 mM Tris buffer, pH 8, 150 mM NaCl, 10% glycerol) to decrease the concentration of non-ionic detergent to less than 0.5%, as recommended by the manufacturer (New England Biolabs). Then 27 μL of diluted lysate was mixed with 1 μL of 0.1 mM SNAP dye (Cat# S9102S, SNAP-Cell 647-SiR), 1 ul 0.1 mM CLIP dye (Cat# S9217S, CLIP-Cell 505) and 1 ul 30 mM DTT and incubated in the dark at room temperature for 1 hour. 10 ul 4X Laemmli sample buffer (Cat# 1610747, BioRad) and 1.5 μL 2-mercaptoethanol (BME) was added to the 30 ul mix, boiled at 95 °C for 5 minutes. After labeling with the SNAP/CLIP substrate, the sample was protected from bright light to avoid photobleaching. 20 μL of the sample was loaded on a gradient stain-free gel (Cat# 4568095, BioRad) to perform SDS-PAGE. The result was imaged with Chemidoc (BioRad) and quantified by image lab software (BioRad).

To label CRY and TIM on resin, ~350 μL cell lysate was mixed with 10 μL pre-washed magnetic HA resin slurry (Cat# 88836, ThermoFisher) and incubated on an end-to-end rotator overnight at 4 °C. The dark sample was covered by foil. Next day, the resin was washed with room temperature 500 μL TBS-T (20 mM Tris buffer, pH 7.6, 150 mM NaCl, 0.05% Tween-20) four times. Then HA resin was resuspended in 27 μL labeling buffer, 1 μL 0.1 mM SNAP dye, 1 μL 0.1 mM CLIP dye and 1 μL 30 mM DTT. The tube was incubated in the dark on an end-to-end rotator at room temperature for one hour. Subsequently, 10 μL 4X Laemelli sample buffer and 1.5 μL BME were added to the 30 μL resin mix. The tube was boiled at 95 °C for 10 minutes to denature and elute CRY and TIM from HA resin. SDS-PAGE and imaging were performed the same as above.

### Clear-Native-Polyacrylamide Electrophoresis

S2 cells were lysed by 2 freeze-thaw cycles to avoid detergent in Native-PAGE samples using the following protocol: An S2 cell pellet from a 5 mL culture was resuspended with 400 μL buffer A (50 mM Tris, pH 8, NaCl 150 mM, 20% glycerol and 1x protease inhibitor; Cat# A32965, ThermoFisher). Then the cell lysate was frozen rapidly in liquid nitrogen and thawed in room temperature water bath. Then the lysate was centrifuged at 20,000 × g for 20 minutes at 4 °C. The supernatant was labeled with SNAP/CLIP dyes (SNAP-Cell 505-Star, Cat# S9103S and CLIP-Cell 505, Cat# S9217S) as above. One part sample was mixed with two parts native sample buffer (Cat#1610738, BioRad) and loaded on the same gradient stain-free gel as above. Clear Native PAGE (CN-PAGE) was performed following the manufacturer instructions, at 150 V for 60 to 90 minutes at room temperature in the dark to prevent photobleaching.

### Expression and purification of sort-tagged H377/H378 CRY variants

All H377/H378 sort-tagged variants of CRY were expressed and purified as has been previously reported (Chandrasekaran et al., 2021). Briefly, the proteins were expressed in CmpX13 cells, a specialized strain of *Escherichia coli* that expresses a riboflavin transporter (Mathes et al., 2009), and were induced with 0.4 mM isopropyl-β-D-1-thiogalactopyranoside (IPTG) at 17 °C. All variants were purified in an identical manner as before using Strep-Tactin resin (Cat# 2-1208-025, IBA).

### Sortylation of CRY variants

The sortylation procedure was carried out as previously reported (1). All reaction mixtures were further purified on an analytical Superdex 200 Size Exclusion Column (10/300 GL, GE Healthcare) in buffer containing 50 mM HEPES (pH 8), 150 mM NaCl, and 10% glycerol (vol/vol). No reductants were added to the gel filtration buffer to avoid reduction of the SORTC-SL nitroxide radical or reduction of the disulfide bond between the spin probe and the peptide.

### ESR spectroscopy measurements (continuous wave and 4P-Double electron electoron resonance; DEER spectroscopy)

All ESR measurements were performed in a similar fashion to what has been previously reported (1). For all ESR experiments, spin-labeled samples were buffer exchanged into deuterated buffer (50 mM HEPES pH 8, 150 mM NaCl) with 25% d8-glycerol utilizing a 50 kDa spin concentrator tube under ambient conditions. The H377A/H378A/L405E/C416N-sort (AAEN-sort) variant was buffer exchanged similarly, but allowed to degas in the glovebox for 1.5 hours under nitrogen in the dark to prevent rapid reoxidation post light exposure. The capillary for this sample was also sealed with epoxy resin prior to removal of the sample from the glovebox. Continuous-wave ESR (cwESR) experiments were performed at X-band (~9.4 GHz) at room temperature with a modulation amplitude of 4 G on a Bruker E500 spectrometer equipped with a super Hi-Q resonator. Samples were illuminated with a blue laser (TECBL-440, 30 mW, 440 nm) for 5-10 s in the resonator cavity. cwESR spectra were taken pre and post irradiation at X-band (~9.4 GHz) prior to DEER measurements. All DEER measurements were carried out at Q-band (~35 GHz) on a Bruker E580 spectrometer equipped with a 10 W solid-state amplifier (150 W equivalent TWTA), 150W RF amplifier, and an arbitrary waveform generator. DEER spectra were measured at 60 K in an EN 5107D2 Cavity with a cryogen-free insert/temperature controller. The measurements were performed using four pulses (π-τ_1_-π_pump_-τ_2_-π-τ_2_-echo) with 16-step phase cycling. The pump and probe pulses (flavin and nitroxide respectively) were separated by 84 MHz (~30 G).

### DEER Spectroscopy Data Analysis

Data analysis for DEER was performed utilizing DD version 7B developed at Vanderbilt University (https://lab.vanderbilt.edu/hustedt-lab/dd/) (Stein et al., 2015) and the Singluar Value Decomposition (SVD) method developed at Cornell University by ACERT (https://denoising.cornell.edu/) (Srivastava and Freed, 2019). In DD, depending on the radical formed, the average distance (〈*R*〉)and width (σ) for either one or two components were fixed while varying the population (in the two-component case) in order to find the best fit in accordance with the docked and undocked states established previously (Chandrasekaran *et al.*, 2021). Unrestrained fits were also performed in DD, where 〈*R*〉 and σ were not fixed, and fits for most variants converged to values previously reported (Chandrasekaran *et al.*, 2021). In addition to the percentage of docked and undocked components, noise corrected error values 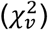 were calculated, 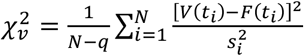, where *V*(*t*) and *F*(*t*) are the experimental and the fit data respectively, *N* is a number of points, *q* is a number of variables changed and *s*_*i*_ is the estimated noise level for the *i*th point. Such values serve as an indication of the fit between the experimental data and the distance reconstruction; values below 2 indicate good fits. The time domain (TD) traces for all H377 and H378 variants were fit directly with linear combinations of the TDs of the WT and L405E/C416N (“EN”) to obtain the relative proportions of the WT (“undocked”) and EN (“undocked”) states. The SVD method was used to determine the distance distributions (P(r)) by obtaining a solution to the Fredholm Equation (K P = S), where P is distance distribution, S is the dipolar signal in the time domain and K is the kernel representing the dipolar interaction between two spins. The SVD method obtains a solution, P(r), by choosing only the large singular values that correspond to the signal and excluding the smaller signaling contributing to the noise.

### UV-Vis Spectroscopy

UV-visible spectra of all alanine variants [in 50 mM HEPES (pH 8), 150 mM NaCl, 10% glycerol (vol/vol)] were taken in a quartz cuvette with a pathlength of 0.2 cm. The spectra were measured using an Agilent 8534 diode-array spectrophotometer with a single reference wavelength set to 800 nm for background correction. All samples were irradiated using a blue laser (TECBL-440, 30 mW, 440 nm, World Star Tech) for 2-5 seconds.

### Molecular Dynamics (MD) simulations

MD simulations were carried out as previously described (Chandrasekaran *et al.*, 2021; Ganguly et al., 2016). The starting structures of the MD simulations were based on the crystal coordinates of full-length Drosophila cryptochrome (PDB ID: 4GU5) similar to the procedure followed by Ganguly et al (Ganguly *et al.*, 2016). dCRY was immersed in an orthorhombic box of rigid TIP3P waters and Na^+^ and Cl^−^ ions were added to produce a neutral physiological salt concentration of 0.15 M. The solvated box was replicated in all three dimensions using periodic boundary conditions and long-range electrostatic interactions were calculated using the particle mesh Ewald (PME) method (Tom Darden, 1993) with a cutoff of 12 Å. Bonds involving hydrogen were constrained by the SHAKE algorithm (Jean-Paul Ryckaert, 1977). After equilibrating the system through several stages that held either pressure or volume constant and varied temperature, (equilibration protocol similar to that used in Ganguly et al PNAS), production trajectories of 200 ns were computed at 298 K in the canonical ensemble (i.e. constant NVT) with a 1 fs time step. A modified Nose-Hoover method in conjunction with Langevin dynamics was employed to maintain constant pressure and temperature during the simulations. All simulations were performed with the AMBER18 software package (Lee et al., 2018), using the pmemd.cuda program (Gotz et al., 2012) and employing the Amber parm99 force field (Cornell et al., 1996). MD simulations were performed with the redox states of the flavin (the neutral state (FADox), the neutral semiquinone (NSQ) state (FADH^•^) and the anionic semiquinone (ASQ) state (FAD—)).

**Figure S1:**
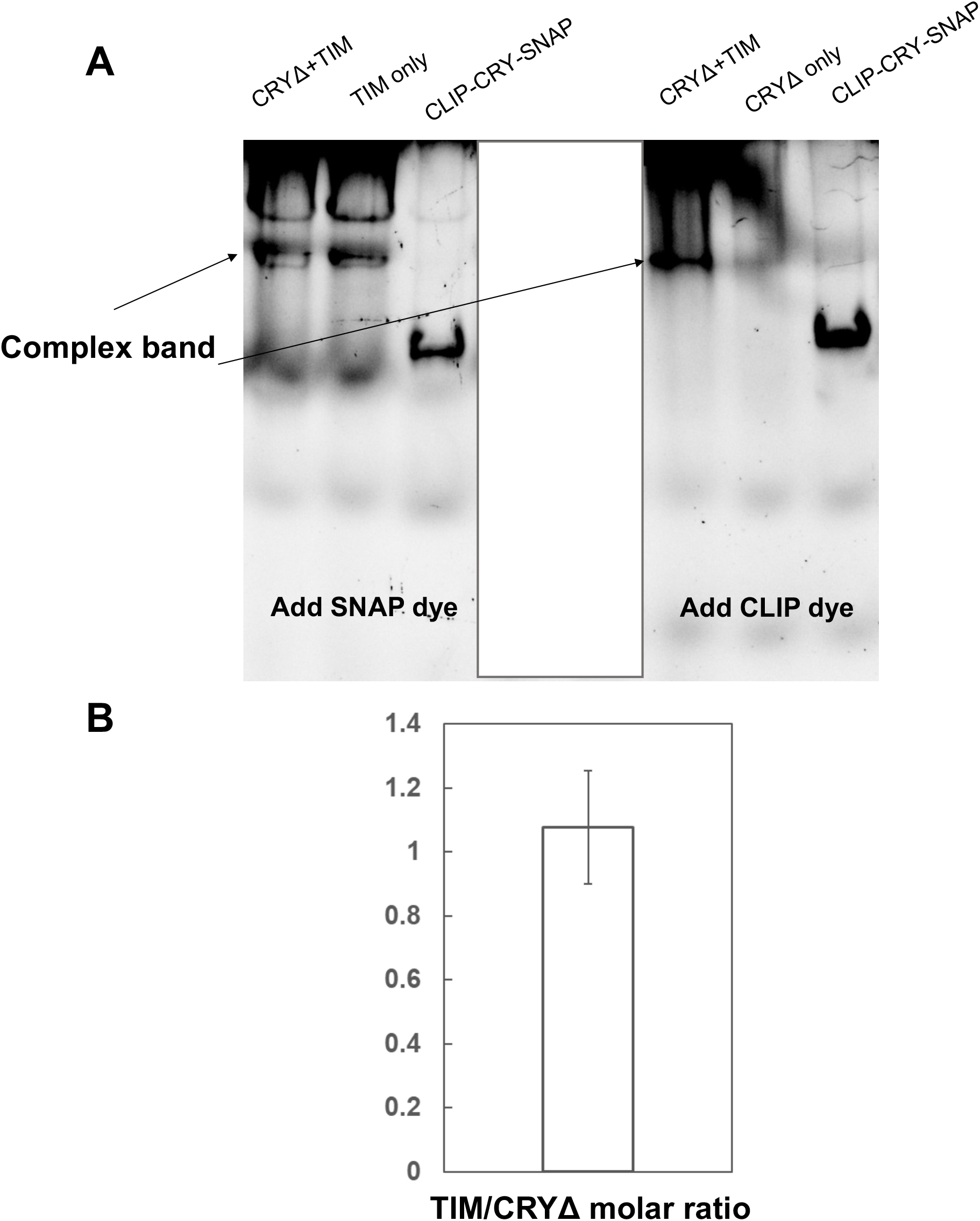
(A) Representative Clear-Native (CN)-PAGE gel of purified CLIP-CRY:TIM-SNAP complexes after dye addition and labeling. Fluorescence detected and imaged by a ChemiDoc (BioRad). Samples on the left were only labeled with SNAP dye and samples on the right were labeled with CLIP dye. Known amount of a CLIP-CRY-SNAP fusion provides the standard for normalization. (B) Quantification of the normalized TIM and CRYΔ fluorescent signal.

**Figure S2:**
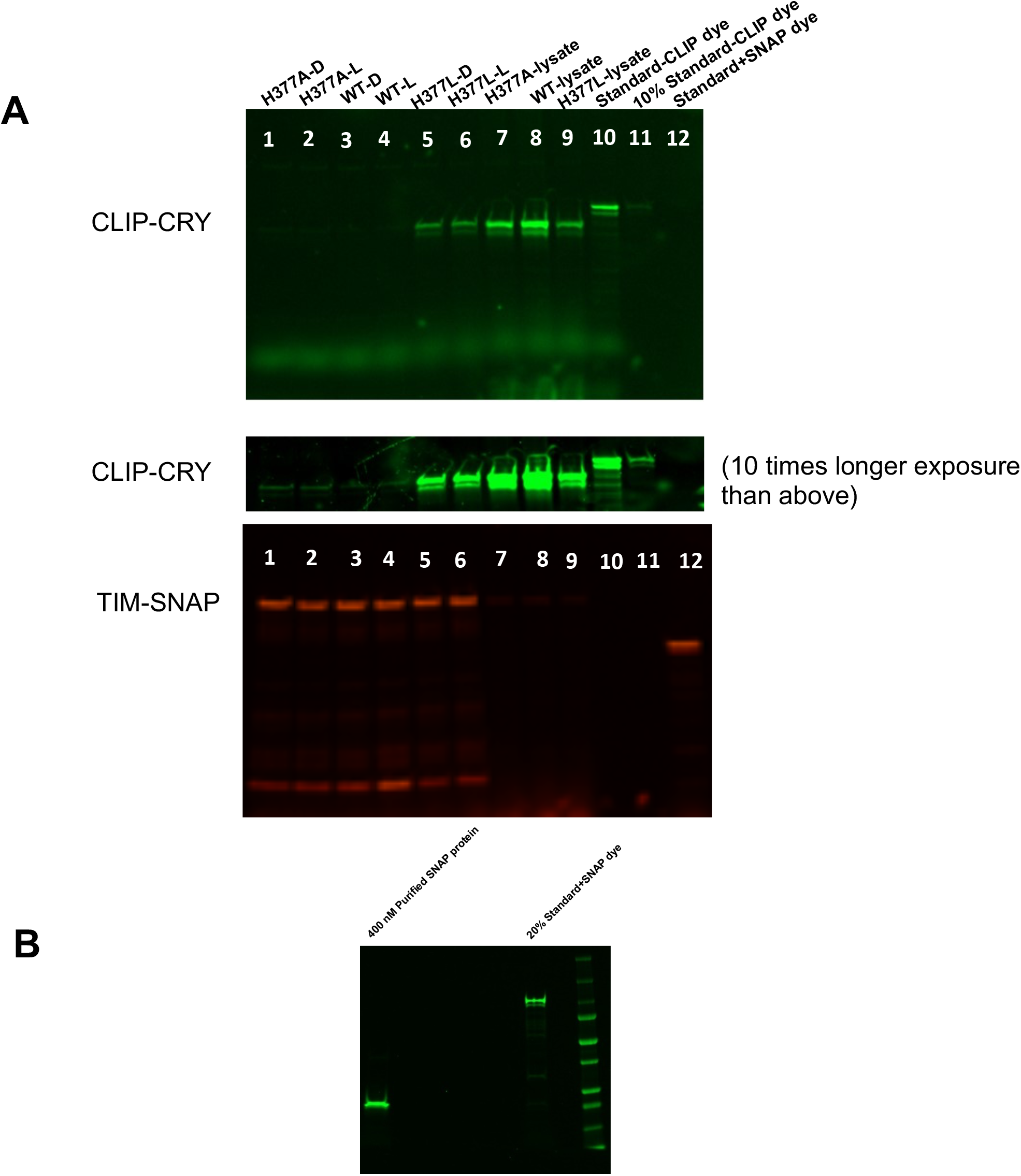
Multiplex imaging of TIM and CRY bands. (A) Lane 1-6 show TIM and CRY SWFTI signals on HA resin, whereas lane 7-9 show the signals in lysate samples. Lane 10-12 exhibit fluorescent signals from the internal standard CLIP-CRY-SNAP. To prevent fluorescence crosstalk, Lane 10 and 11 have the standard only mixed with CLIP dye, whereas lane 12 contains the standard with only SNAP dye. In lane 11 the standard was diluted to 10%. (B) The quantification of standard by purified SNAP proteins. The standard was diluted to 20% and mixed with SNAP dye.

**Figure S3:**
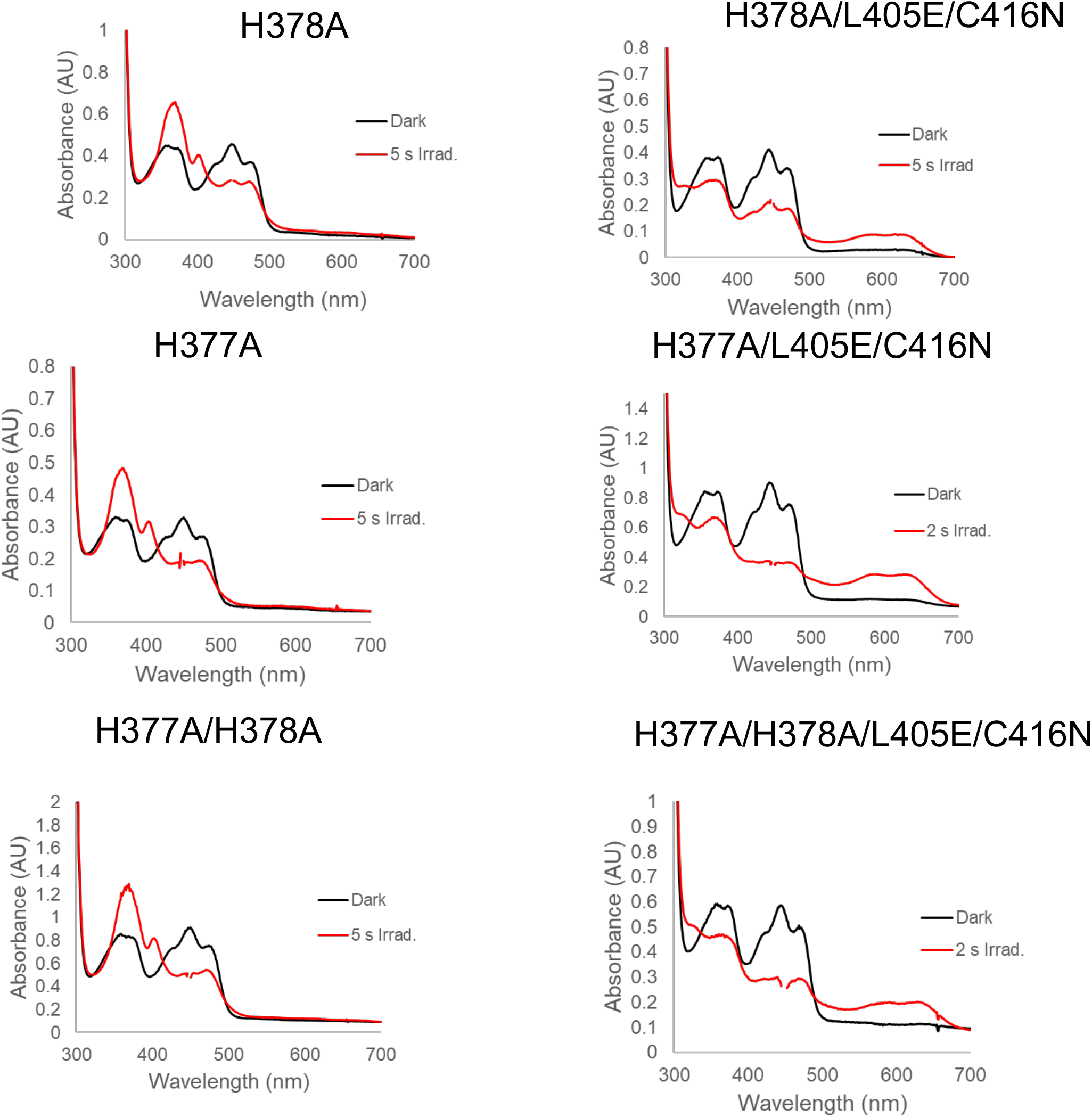
UV-Vis spectra of His377 and His378 alanine variants in the dark and after 2-5 sec of blue light exposure at 440 nm. The left column are all variants in the WT background, while the right column is those in the L405E/C416N background. All WT background variants produce the ASQ with a characteristic absorbance at 403 nm, while the L405E/C416N background variants produce the NSQ (broadband feature ~550-650 nm). Gaps in spectra are at the excitation laser wavelength; Time of irradiation for full reduction varies between 2-5 second depending on laser alignment with respect to the cuvette.

**Figure S4:**
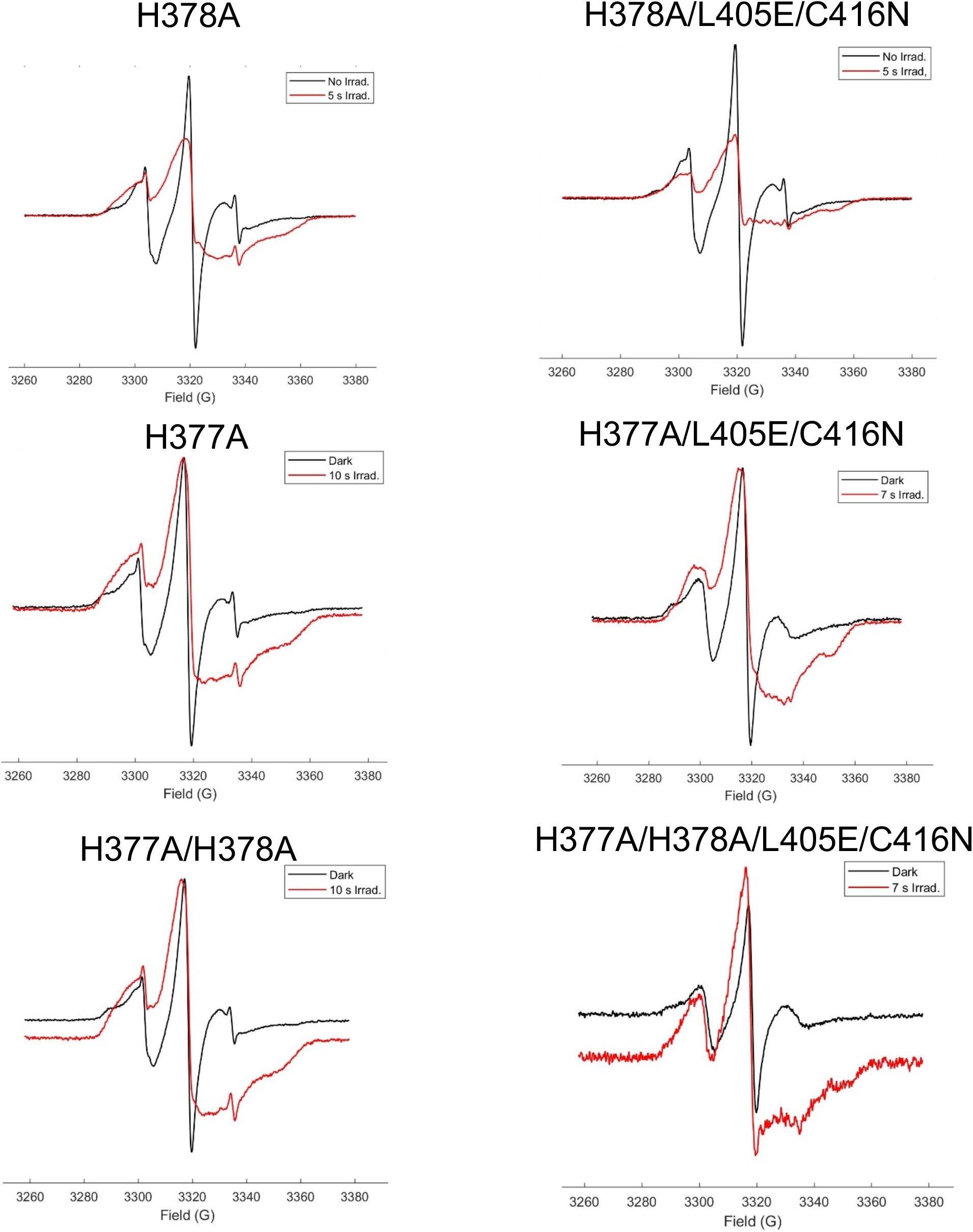
cwESR spectra of all nitroxide-labeled dCRY variants in the dark and post light exposure. All spectra were recorded at X-band in deuterated buffer. Broadened features in light-state spectra reflect overlapping flavin and nitroxide features.

**Figure S5:**
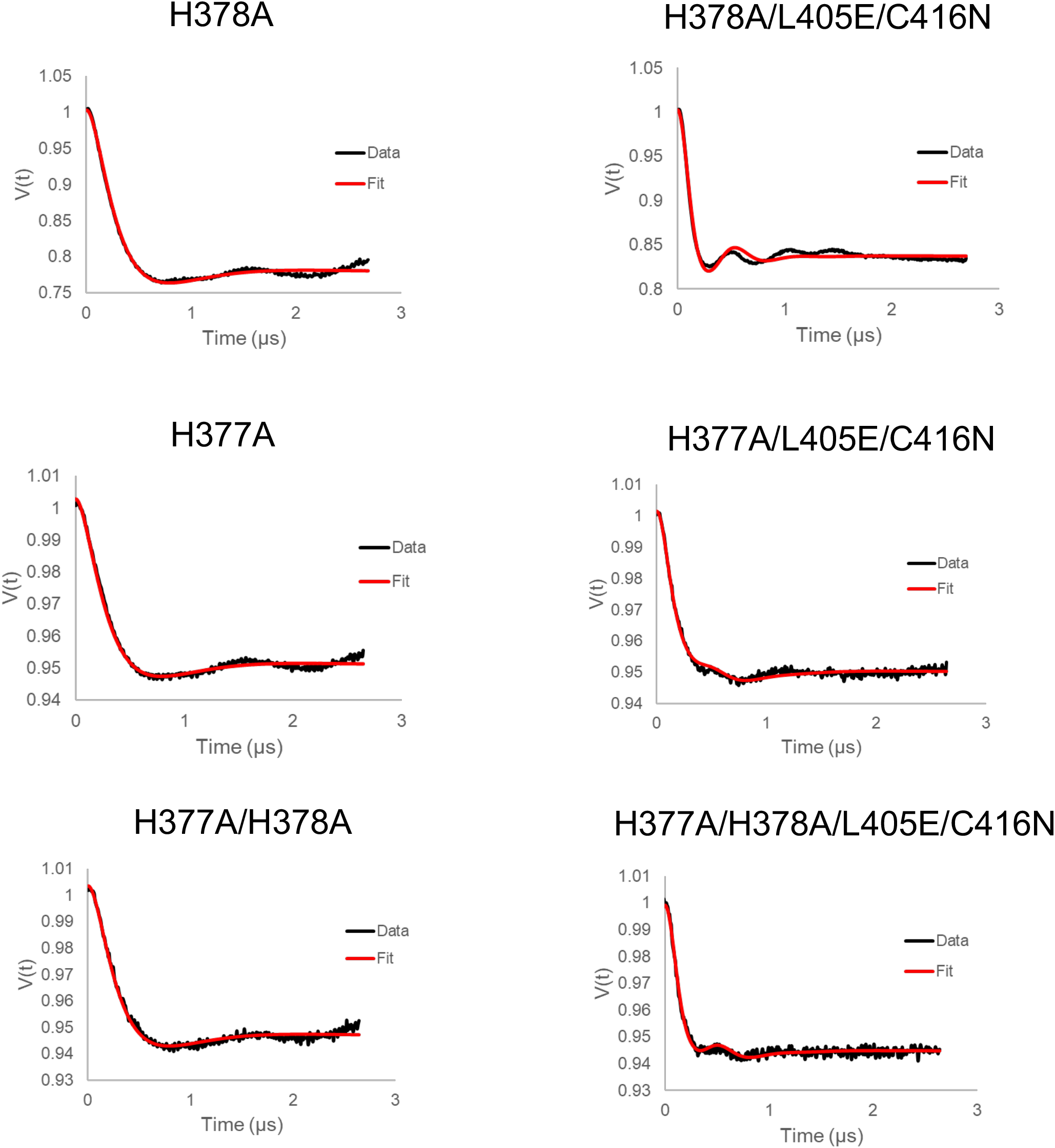
Time domain traces of all alanine variants and their respective fits carried out based on the two-state model previously established by using DD. All ASQ forming variants (left) were fit with one component, while all NSQ forming variants (right) were fit using two components. Fittings were carried out as detailed in the **Methods**.

**Figure S6:**
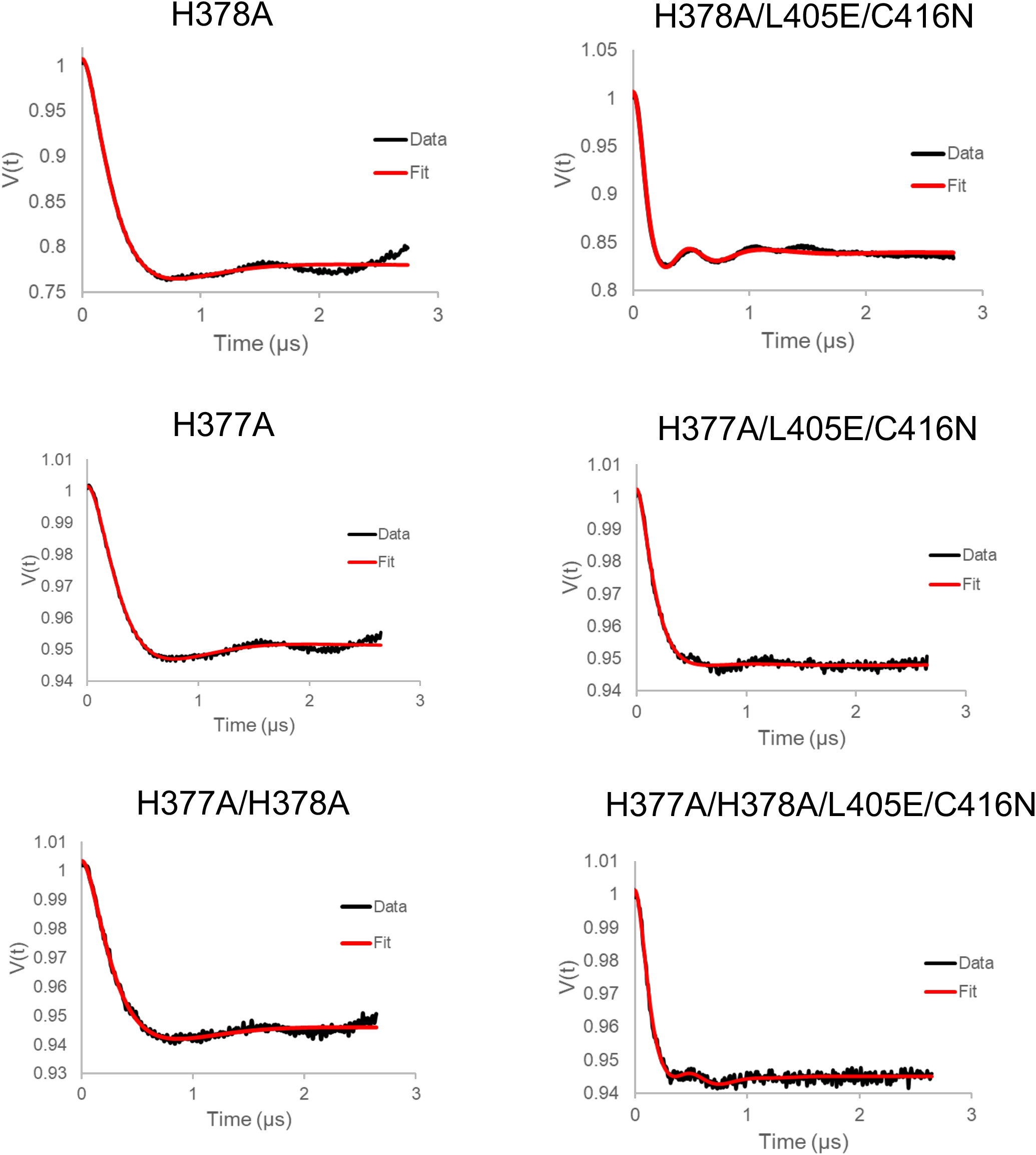
Unrestrained fittings by DD to experimentally obtained time domain data.

**Table S2:** Unrestrained DD fitting statistics. Residual error χv^2^ values are determined by DD and indicate the robustness of the fit to the data. Values below 2 indicate a good fit. H377A, H378A, and AA are all WT-like and were fit best using one-component assuming complete undocking (*). The relatively high error (**) reflects the small amplitude and broad features of this component.

| Variant | Radical | % Undocked | $\chi_v^2$ (error in fit) |
| --- | --- | --- | --- |
| H377A | ASQ | 100* | 2.6** |
| H378A | ASQ | 100* | 5.4** |
| H377A/H378A | ASQ | 100* | 1.2 |
| H377A/L405E/C416N | NSQ | $19 \pm 97^{***}$ | 1.2 |
| H378A/L405E/C416N | NSQ | $21 \pm 0.5$ | 6.3** |
| H377A/H378A/L405E/C416N | NSQ | $64 \pm 7$ | 0.8 |

**Figure S7:**
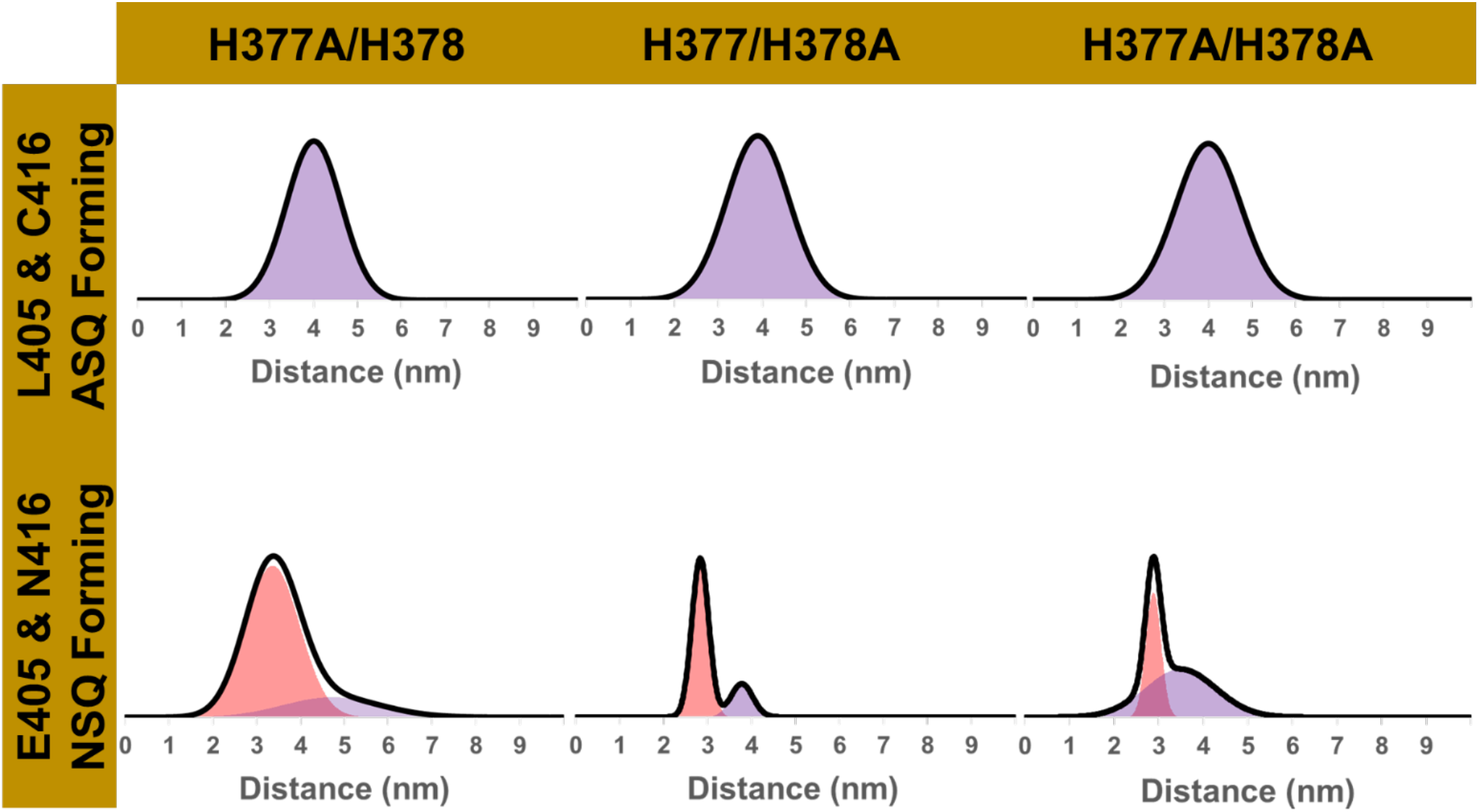
Distance distributions of all alanine variants produced from unrestrained fits by DD.

**Figure S8:**
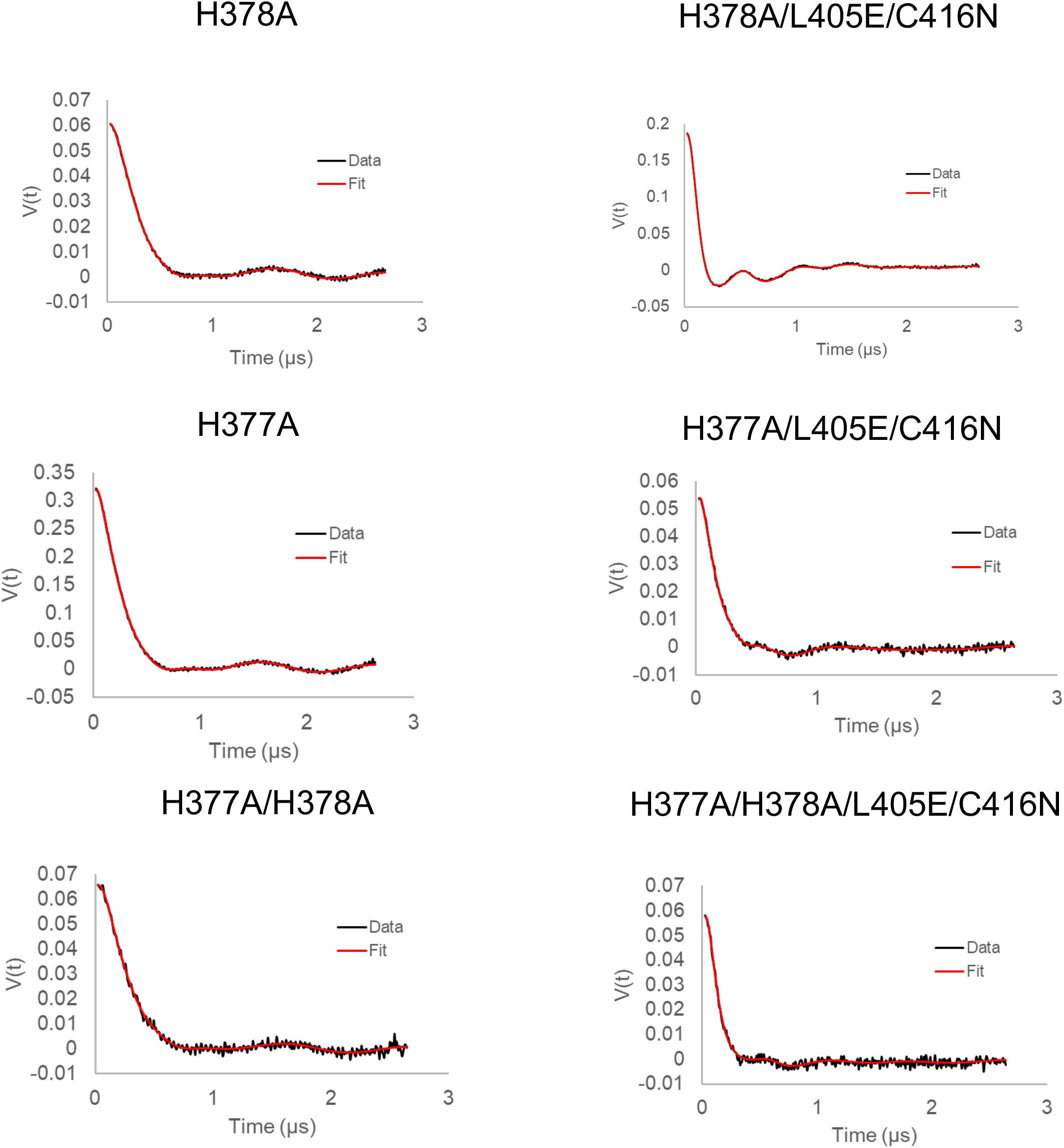
Time domain fittings of all alanine variants using the SVD method as compared to the experimentally obtained data.

**Figure S9:**
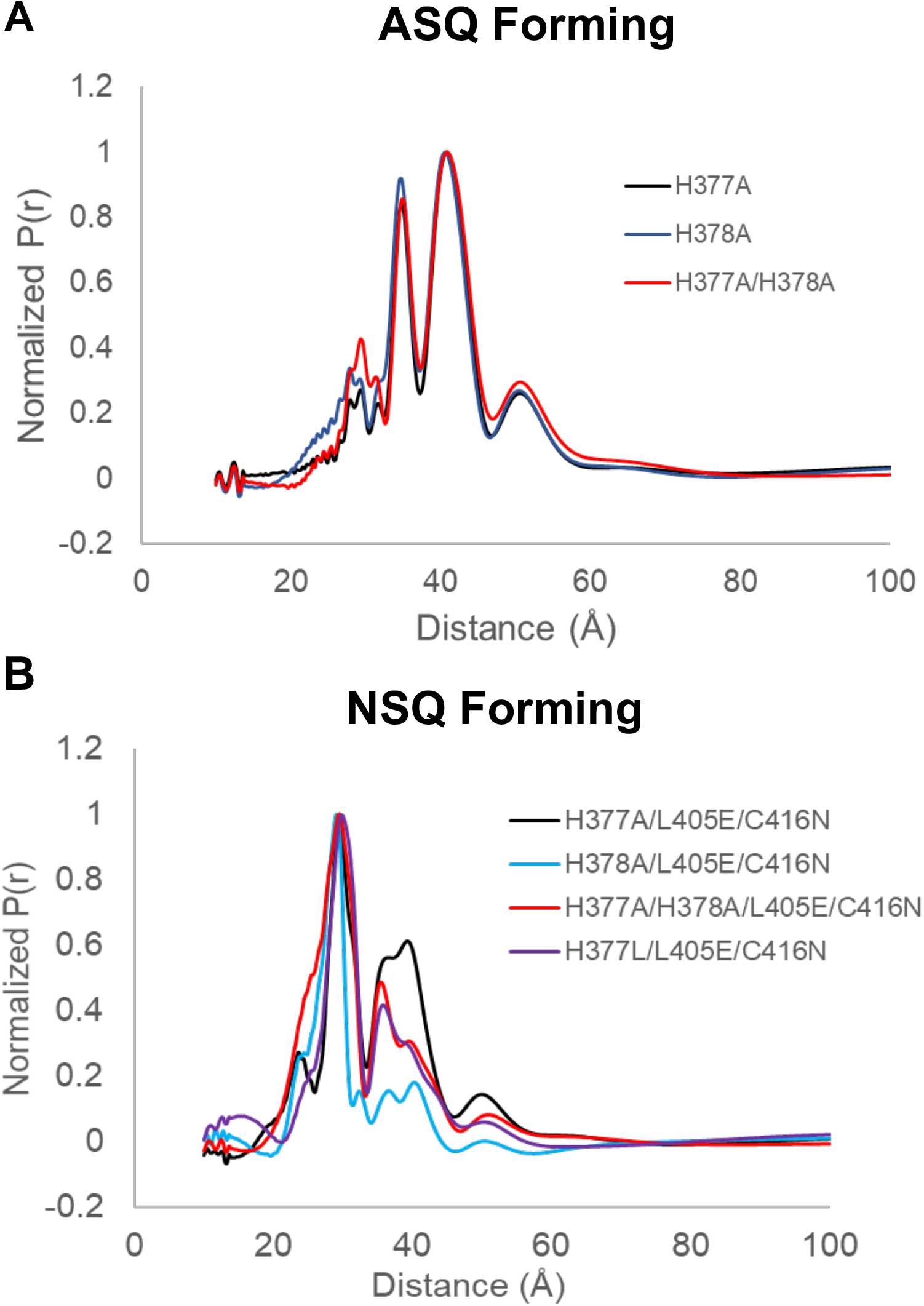
Distance distributions obtained using the SVD method for (a) ASQ and (b) NSQ forming variants.

**Table S3:** Percent undocked state for Ala variants using linear combination fit. Values determined by fitting linear combinations of WT and EN time domain traces to time domain data of all variants.

| Variant | Radical | % Undocked |
| --- | --- | --- |
| H377A | ASQ | 100 |
| H378A | ASQ | 100 |
| H377A/H378A | ASQ | 100 |
| H377A/L405E/C416N | NSQ | 58 |
| H378A/L405E/C416N | NSQ | 17 |
| H377A/H378A/L405E/C416N | NSQ | 34 |

**Figure S10:**
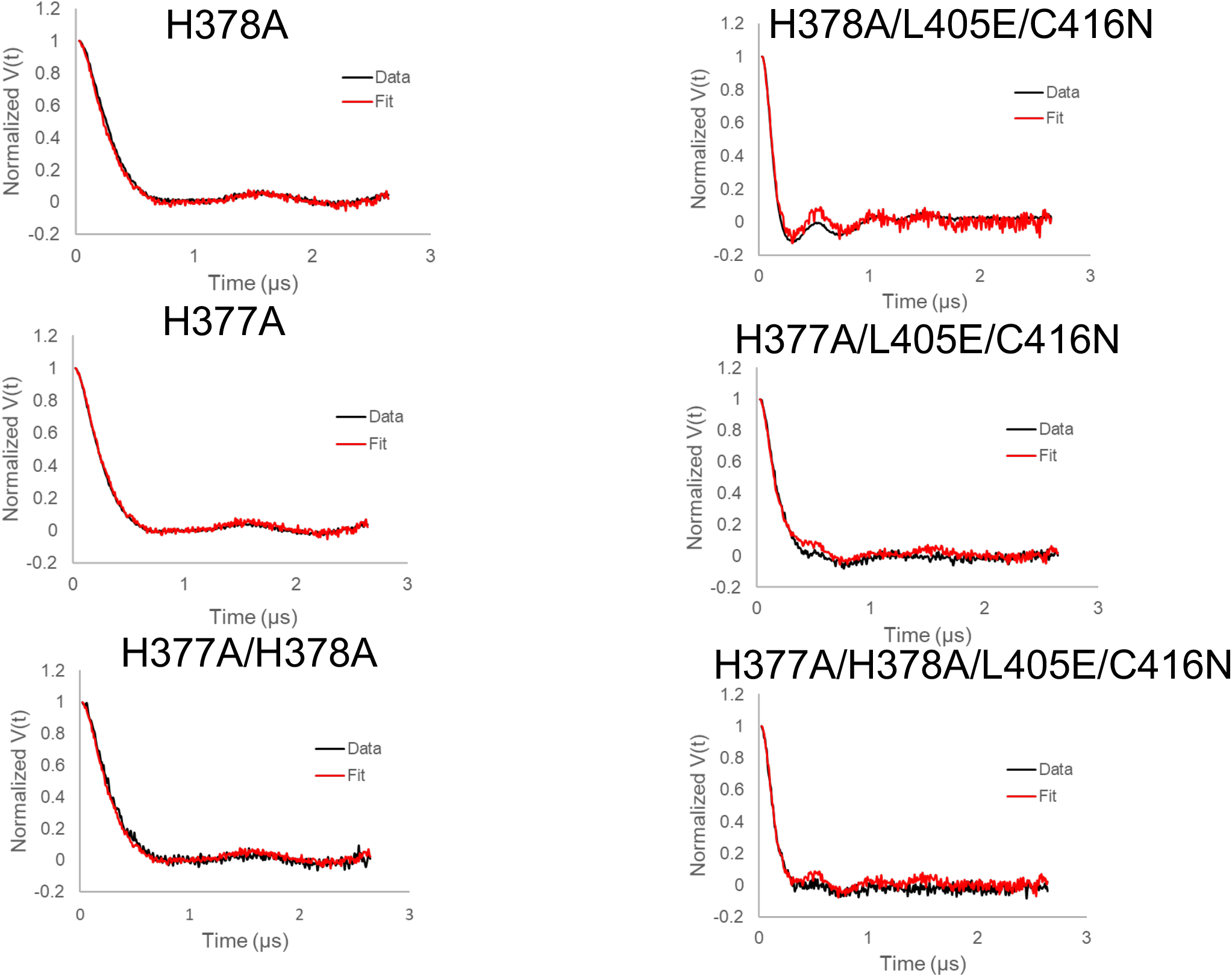
Time domain fits by linear combinations of All the WT and EN time domain traces previously determined. The best fit was determined by varying the proportions of each and validating the fit by visual inspection. The percentages obtained by the fitted linear combinations agree well with those obtained by the restrained fits obtained in DD (Table 1).

**Figure S11:**
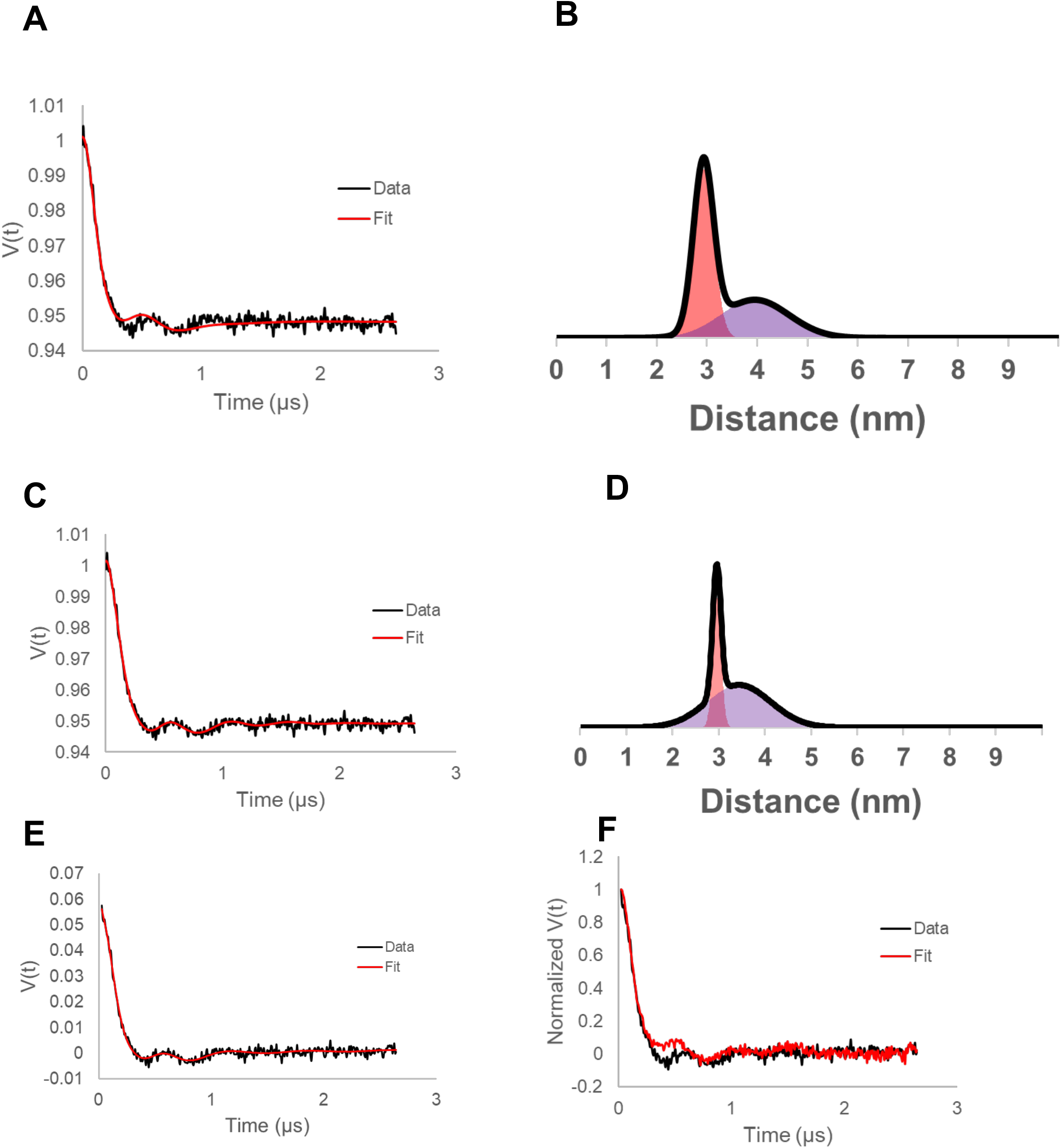
Time domain reconstruction (A) and distance profile (B) restricted to the two-state model in DD (% Undocked = 43%; Χv^2^ (restrained) = 0.39264). Time domain reconstruction (C) and distance profile (D) unrestrained in DD (% Undocked = 72%; Χv^2^ (unrestrained) = 0.27531). (E) Time domain fitting from SVD. (F) Linear combination fit using WT and EN time domain traces (% WT-like (“Undocked”) = 42%).

